# GPR15LG binds CXCR4 and synergistically modulates CXCL12-induced cell signaling and migration

**DOI:** 10.1101/2025.01.13.632759

**Authors:** Dan Albers, Sofya Novikova, Julio Vieyto-Nuñez, Yasser Almeida-Hernández, Chiara Pastorio, Florian Klassen, Dana Weiss, Pascal von Maltitz, Janeni Jaikishan, Moumita Datta, Hassan Jumaa, Billy Jebaraj, Stephan Stilgenbauer, Manish Kumar, Palash Chandra Maity, Christian Buske, Ulrich Stifel, Julia Zinngrebe, Pamela Fischer-Posovszky, Andy Chevigné, Frank Kirchhoff, Elsa Sanchez-Garcia, Jan Münch, Mirja Harms

## Abstract

GPR15LG, a chemokine-like ligand for the G-protein coupled receptor 15 (GPR15), is abundantly expressed in the gastrointestinal mucosa and inflamed skin. Emerging evidence suggests its involvement in inflammatory disorders and cancers. This study investigates the effects of GPR15LG on the signaling and downstream functions of C-X-C chemokine receptor type 4 (CXCR4), which plays a critical role in immune cell trafficking and cancer metastasis. The results demonstrate that GPR15LG binds to the orthosteric site of CXCR4, modulating downstream signaling in a context-dependent manner. Specifically, GPR15LG enhances CXCL12-mediated CXCR4 signaling synergistically, promoting wound healing and cell migration across various cell types, including CD4+ T cells and cancer cells. These findings underscore the role of GPR15LG in inflammation and metastasis, offering potential therapeutic avenues for CXCR4-mediated diseases.

**Teaser:** GPR15LG binds CXCR4 thereby modulating CXCL12/CXCR4 signaling and immune and cancer cell trafficking.

## 1. Introduction

The G protein-coupled receptor 15 ligand (GPR15LG), also known as C10orf99 or AP-57, is a highly cationic protein increasingly recognized for its role in modulating epithelial inflammation and immune responses ^1,2^. GPR15LG is implicated in a variety of biological processes, including antimicrobial defense, chemotaxis, epithelial homeostasis, and the regulation of both innate and adaptive immune responses ^3^. Notably, its expression is highest in epithelial tissues, particularly in the gastrointestinal mucosa and inflamed skin. This underscores its potential role in maintaining epithelial integrity and mediating immune defenses in these environments ^2^.

Recently, GPR15LG was identified as the chemokine-like ligand of the G protein-coupled receptor GPR15, which mediates lymphocyte homing to the gut and skin ^1,2,4,5^. In healthy tissues, GPR15LG has been reported to play a homeostatic role, contributing to the maintenance of epithelial barriers and the modulation of local immune responses ^1,2^. Dysregulation of GPR15LG has been associated with various infectious and inflammatory disorders. For example, GPR15LG expression is significantly upregulated in chronic inflammatory skin diseases, such as psoriasis and atopic dermatitis, as well as the wound borders of chronic ulcers ^6–9^. However, the precise role of GPR15LG in these processes is unclear. Previous studies yielded opposing data with some reports suggesting that it may exacerbate inflammatory processes ^7,10,11^, while others propose anti-inflammatory or tissue-protective properties ^12,13^. Thus, GPR15LG exerts complex effects that may vary between different cell and tissue types and stages of inflammation.

In addition to its role in inflammatory diseases, emerging evidence indicates an involvement of GPR15LG and its receptors in cancer biology, particularly in colorectal cancer ^14,15^. Specifically, it has been reported that GPR15LG exerts tumor-suppressive effects by inhibiting cell proliferation and promoting cell cycle arrest ^14^. Others studies showed, however, increased infiltration of GPR15+ Tregs into the tumor microenvironment and suppression of immune function ^14,16^. These seemingly contradictory findings reflect the complex and multifaceted nature of GPR15LG’s function, which may depend on the repertoire of interacting receptors and targeted signaling pathways.

Thus, many questions regarding GPR15LG function and potential interactions with receptors other than GPR15 remain. Recently, we found that GPR15LG competes with an antibody specific for CXCR4 ^17^, a GPCR involved in immune cell trafficking, homeostasis and tissue repair. The cognate ligand for CXCR4 is CXCL12, also known as stromal derived factor 1 (SDF-1) ^18^. Both, CXCR4 and CXCL12, are frequently upregulated during inflammation and contribute to disease progression, especially in chronic conditions ^19,20^. In addition, CXCR4 is upregulated in many types of cancer, especially on cancer stem cells, and contributes to their survival, proliferation and migration ^18,19,21,22^. Whether or not GPR15LG modulates CXCR4-CXCL12 signaling and immune or cancer cell migration remains to be determined.

In addition to CXCR4, CXCL12 also binds to the atypical chemokine receptor 3 (ACKR3) inducing β-arrestin-2 recruitment and receptor internalization ^23^. ACKR3, also known as CXCR7, functions primarily as a scavenger receptor for CXCL12, regulating its extracellular concentrations and modulating CXCL12-CXCR4-mediated signaling ^24^. Similar to CXCR4, ACKR3 has implications for organ development, cell migration, and positioning during embryogenesis. In adulthood, it contributes to tissue homeostasis and is implicated in various cancers, autoimmune disorders, and cardiovascular conditions ^24,25^.

Here, we examined the interactions of GPR15LG with CXCR4 and ACKR3 and their consequences on downstream signaling as well as immune and cancer cell trafficking.

## 2. Results

### 2.1. GPR15LG interacts with CXCR4 and inhibits CXCR4-tropic HIV-1

We have previously shown that GPR15LG competes with the CXCR4-specific antibody clone 12G5 for binding to CXCR4 ^17^. To further define this interaction, SupT1 cells were exposed to two monoclonal CXCR4 antibodies in the presence of serially diluted GPR15LG, CXCL12, or AMD3100, a small molecule CXCR4 antagonist. The antibody clone 12G5 binds close to the CXCR4 binding pocket ^26,27^ and clone 1D9 to the N-terminus of CXCR4 ^28^. As expected, AMD3100, which binds orthosterically to CXCR4, replaced only 12G5 (IC_50_ = 383 ± 40 nM) but not 1D9 ^28^. In contrast, CXCL12 that interacts with CXCR4 via the N-terminus and the binding groove ^29^ competed with both antibodies (**Figure 1a**). GPR15LG also competed with both CXCR4 antibodies (IC_50_ = 1.3 ± 0.1 µM for 12G5 and ∼ 12.1 ± 4.4 µM for 1D9), suggesting a similar binding mode to the receptor as CXCL12.

**Figure 1.**
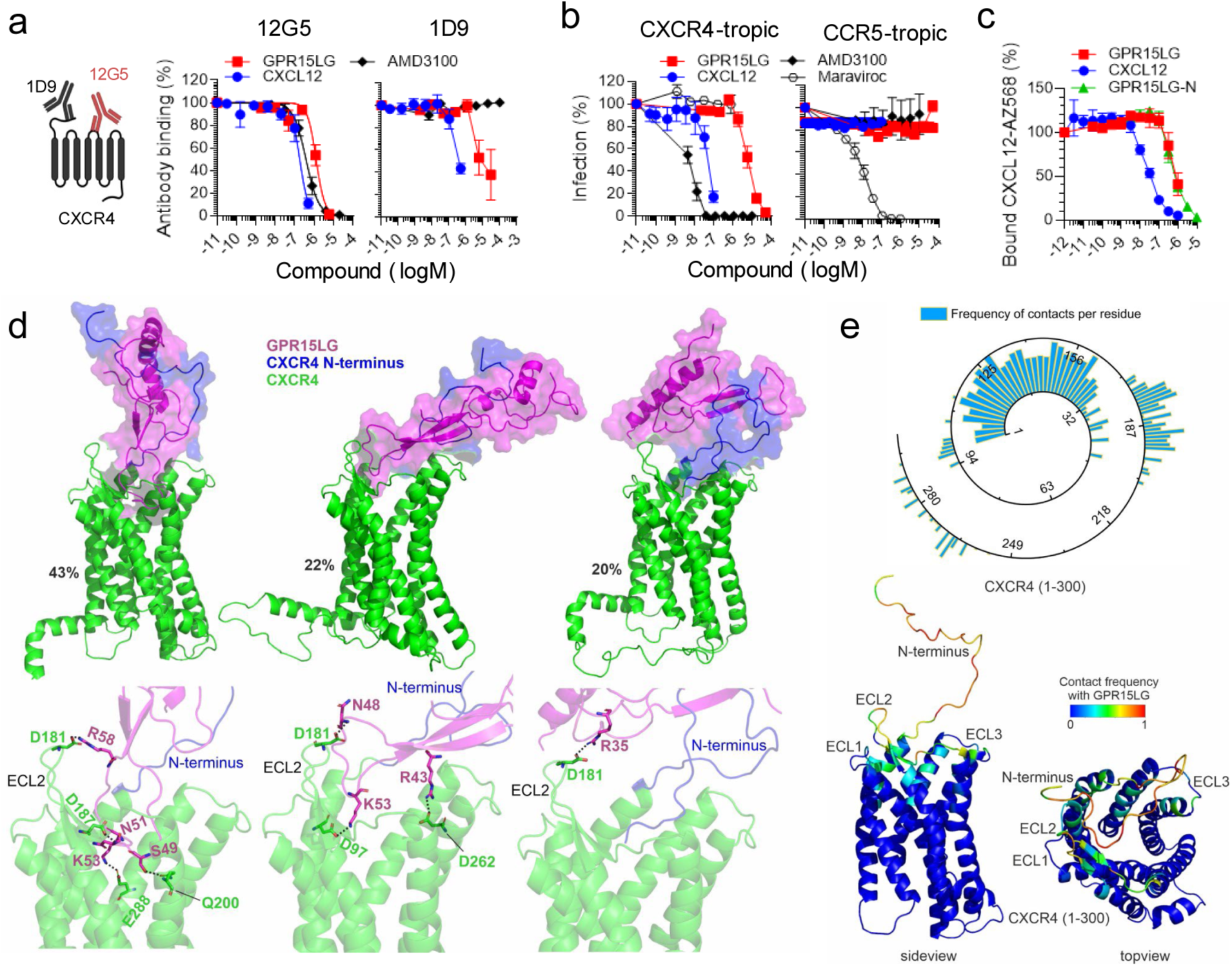
GPR15LG interacts with the orthosteric pocket of CXCR4. **(a)** GPR15LG competes with CXCR4 antibodies for binding. SupT1 cells were incubated with fluorescently labeled antibodies (clone 12G5, targeting CXCR4’s extracellular loops 1 and 2, and clone 1D9, targeting the CXCR4 N-terminus) in the presence or absence of GPR15LG. Binding was analyzed by flow cytometry. **(b)** Binding competition of CXCL12-AZ568 with GPR15LG was determined on 293T cells stably expressing CXCR4 fused to a Nanoluciferase and monitored by NanoBRET. **(c)** GPR15LG inhibits CXCR4-tropic HIV-1 infection, but not CCR5-tropic HIV-1 infection. TZM-bl cells were inoculated with either CXCR4- or CCR5-tropic HIV-1 strains in the presence of GPR15LG. Infection was measured 3 days post-infection by β-galactosidase assay. **(a-c)** Data are shown as mean ± SEM, *n = 3*. **(d)** Results from GaMD simulations of GPR15LG with CXCR4. Computationally predicted binding modes of full-length GPR15LG (magenta) and CXCR4 (green, except for N-terminus in blue). The top three most populated clusters of GPR15LG are shown with special focus on the hydrogen bond interactions at the binding pocket of CXCR4. **(e)** Contact frequency between residues of GPR15LG and CXCR4. The frequency of interactions (scaled 0–1) is mapped onto the CXCR4 structure, highlighting key regions such as extracellular loops 1 (ECL1: 100–104), ECL2 (174–192), and ECL3 (267–273).

HIV-1 uses CXCR4 or CCR5 as coreceptors for entry into target cells, and ligands binding to these receptors inhibit HIV-1 infection ^30^. To examine whether GPR15LG binding to CXCR4 interferes with HIV-1 infection, we incubated TZM-bl reporter cells with GPR15LG, CXCL12, AMD3100, and the CCR5 antagonist Maraviroc and subsequently infected them with CXCR4- or CCR5-tropic HIV-1. As expected, CXCL12 and AMD3100 specifically inhibited CXCR4-tropic HIV-1 infection, with IC_50_ values of 47.9 ± 8 nM and 5.3 ± 0.4 nM, respectively (**Figure 1b**), while Maraviroc blocked CCR5-tropic HIV-1 infection ^31–33^. GPR15LG showed dose-dependent inhibition of CXCR4-tropic HIV-1 infection (IC_50_ = 5.9 ± 0.54 µM) but had no effect on CCR5-tropic HIV-1, further confirming its specificity for CXCR4.

We next tested for the displacement of labelled CXCL12 from CXCR4. Unlabeled CXCL12, which was used as a control, led to a displacement with an IC_50_ value of 36.7 ± 5 nM (**Figure 1c**). GPR15LG dose-dependently competed with labelled CXCL12 with an IC_50_ value of 829 ± 139 nM, confirming CXCR4 interaction.

### 2.2. **C**omputational model of GPR15LG binding modes with CXCR4

To gain deeper insights into the interactions between GPR15LG and CXCR4, we conducted enhanced sampling simulations using Gaussian accelerated molecular dynamics (GaMD). We employed an unbiased approach, positioning GPR15LG 100 Å from the receptor’s center of mass in three distinct orientations (**Figure S1**). For each initial pose, three replicas (400 ns each) of the production MD simulations were carried out and the analysis focused on the final 200 ns of each replica after binding of GPR15LG to the receptor. **Figure 1d** displays three different binding modes of GPR15LG to CXCR4, obtained from clustering analyses of the GaMD trajectories. The most populated cluster of structures (43% of the sampled conformations) displays the main binding mode. In this mode, GPR15LG primarily interacts with the N-terminal region of CXCR4 and also inserts itself into the orthosteric pocket of CXCR4. Notably, GPR15LG establishes consistent interactions with the second extracellular loop (ECL2, amino acids 174-192), which connects TM3 to TM4 and is critical for ligand binding in many GPCRs, including CXCR4. This binding mode aligns with GPR15LG’s competitive interaction with the antibodies 1D9 and 12G5, which targets the N-terminal and ECL2 regions, respectively. The second and third most populated binding modes (22% and 20%, respectively) involve interactions more focused on the ECL2 region, and less penetration of GPR15LG into the orthosteric pocket of CXCR4. The importance of the N-terminus region of CXCR4 in the initial recognition of GPR15LG is common for all binding modes, also guiding GPR15LG toward the receptor’s orthosteric pocket in the most populated binding pose. This binding mode emphasizes the role of the N-terminus of the receptor in complex formation, as previously reported for CXCR4-CXCL12 interactions ^29^.

Hydrogen bond interactions established between GPR15LG and CXCR4, as observed along the GaMD simulations, involve key contacts between GPR15LG’s residues R35, R38, R43, N48, S49, N51, K53 and R58 and CXCR4’s residues D97, D181, D187, D193, Q200, D262, E268, E277, and E288 (**Figure 1e, Figure S2**). The first five residues from GPR15LG are located in the 25-41 segment (GPR15LG-N), which suggests a crucial role of the N-terminus in binding to the receptor, as confirmed by displacement experiments (**Figure 1c**). Besides, positively charged residues in GPR15LG - such as R38, R43, K53, and R58 - are conserved across different GPR15LG sequences from various species, highlighting their functional importance (**Figure S3**). These electrostatic interactions between charged residues of GPR15LG and CXCR4 play a crucial role in stabilizing the ligand-receptor complex as previously proposed ^34,35^. For example, derivatives of the EPI-X4 peptide contain Lys and Arg residues critical for binding to CXCR4 ^34,35^. A previous study combining cryo-EM and MD modeling of the CXCR4-CXCL12 complex ^29^ reported a binding mode similar to the most populated CXCR4-GPR15LG complex. Consistently, residues D97, D262, and E288 of CXCR4 establish conserved hydrogen bonds with residues K1, R8, V3, and S4 from the N-terminus of CXCL12.

Our simulations show that GPR15LG interacts with CXCR4 through three main binding modes, mainly targeting the N-terminus in its way to accessing ECL2, and the orthosteric pocket. These interactions enable GPR15LG to competitively inhibit CXCL12 binding, where electrostatic forces play a key role in complex stability. The similarity of GPR15LG’s binding to CXCR4-CXCL12 complexes highlights its potential as a modulator of CXCR4 signaling, potentially influencing immune and cancer cell regulation.

### 2.3 Computational model of GPR15LG binding to ACKR3

CXCL12 also interacts with the atypical chemokine receptor 3 (ACKR3), a decoy receptor that modulates chemokine availability and signaling without G-protein signaling ^36^. To explore whether GPR15LG could interact with ACKR3, we again used GaMD simulations (**Figure 2**). Our analysis of the first and second simulation clusters indicates that GPR15LG primarily interacts with the N-terminus of ACKR3, with only partial interactions with residues at the top of the binding pocket, such as E207, D275 and E290 (**Figure 2a, Figure S2**). Notably, the N-terminus of ACKR3 is about 10 amino acids longer than that of CXCR4, and its folding appears to hinder the entry of GPR15LG into the ligand-binding pocket. This extended N-terminus likely acts as a steric hindrance, limiting a deeper engagement of GPR15LG with the binding pocket of ACKR3, a behavior not observed with CXCR4. To test this assumption experimentally, we performed competition assays with labeled CXCL12 and found that GPR15LG does not compete with CXCL12 binding to ACKR3 (**Figure 2b**).

**Figure 2.**
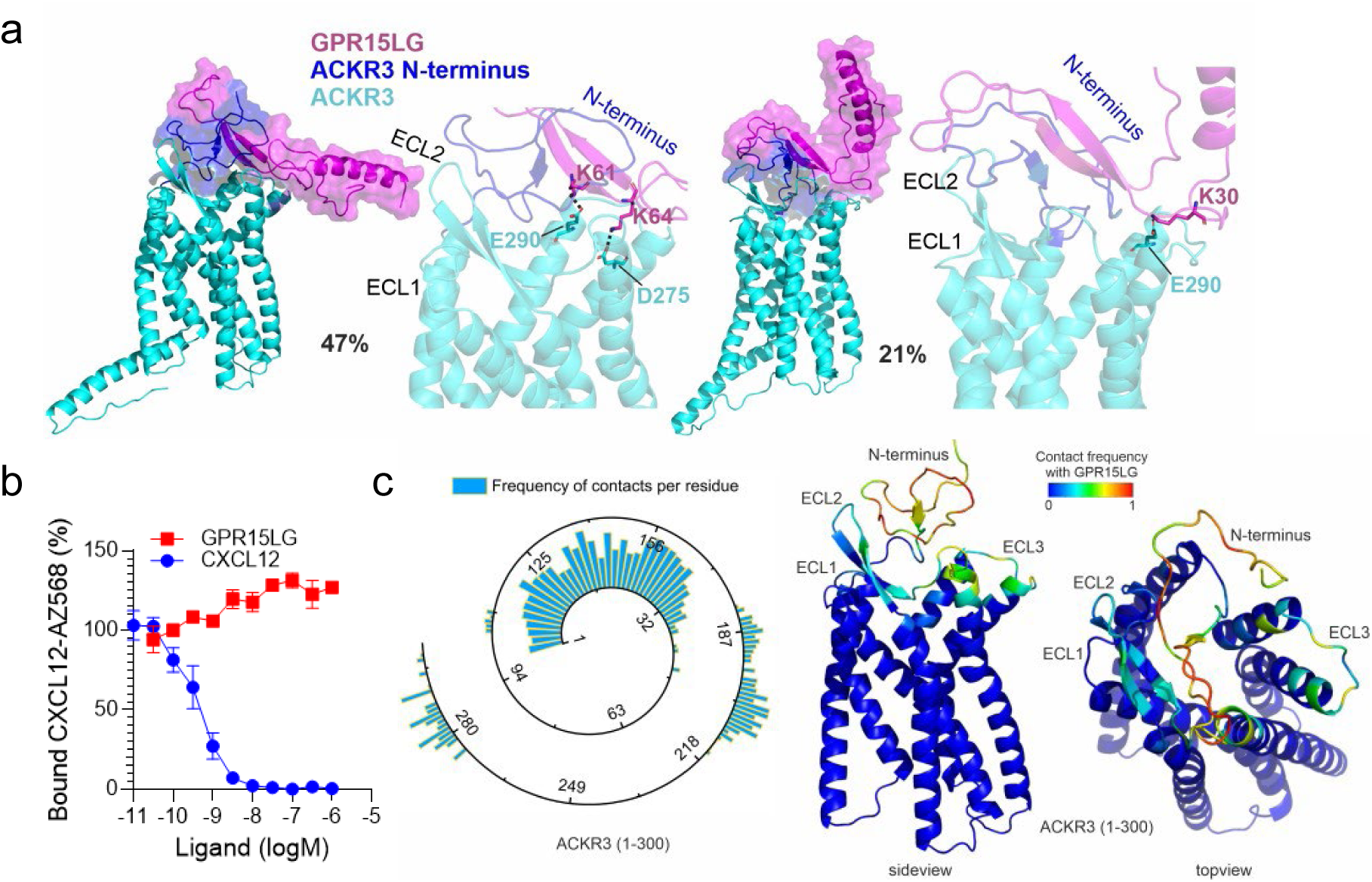
GPR15LG remains outside the binding pocket of ACKR3. a) Results from the GaMD simulations of GPR15LG with ACKR3. Computationally predicted binding modes of the full-length GPR15LG (magenta) with ACKR3 (cyan, except for the N-terminus in blue). The two most populated clusters of GPR15LG are shown with special focus on the hydrogen bond interactions at the binding pocket of ACKR3. b) Binding competition of CXCL12-AZ568 with GPR15LG was determined on HEK293T cells stably expressing ACKR3 fused to a Nanoluciferase and monitored by NanoBRET. c) Frequency of contacts between residues of GPR15LG and ACKR3. The frequency of contacts (scaled 0–1) between GPR15LG and ACKR3 (amino acids 1–300) is mapped onto the ACKR3 structure, highlighting key binding regions.

GPR15LG exhibits higher frequency of contacts (within 5 Å between heavy atoms) with the ECL motifs at the CXCR4 binding pocket compared to ACKR3 (**Figures 1e, 2c**). For example, the sequence segment around D187, which corresponds to ECL2 (residues 174–192) in CXCR4, shows a greater population of contacts per residue than the equivalent region in ACKR3. This finding is further supported by the average α-carbon distances between GPR15LG and the receptors (**Figure S4**). While both receptors share similar contact areas with GPR15LG, CXCR4 features a broader range of interactions and closer proximity (<30 Å) at the 90–120, 180–210, and 250–300 sequence segments compared to ACKR3. These regions are critical for GPCR function as they include key structural motifs of the receptors, such as the ECLs and residues of the binding pocket. The proximity to GPR15LG (<20 Å α-carbon distance) and the broader interaction range in the case of CXCR4 suggest a higher likelihood of interaction and affinity, potentially explaining the more effective engagement of GPR15LG with CXCR4. Additionally, the differences in N-terminal length and the resulting steric effects observed in the case of ACKR3 highlight the distinct ligand recognition and binding dynamics between the two receptors.

### 2.2. GPR15LG is an agonist for GPR15, but not CXCR4

Our results suggest that GPR15LG is not only an agonist for GPR15 but also interacts with CXCR4. In addition, modeling data suggest an interaction with ACKR3, which was, however, not supported by the CXCL12 competition assay. Typically, GPCR activation, including GPR15 and CXCR4, triggers G-protein dissociation, activating downstream pathways like Akt and Erk phosphorylation, along with calcium release ^18,37^. Signaling is terminated by receptor phosphorylation, β-arrestin recruitment, and internalization ^18,37^. In contrast, ACKR3 is an atypical chemokine receptor, interacting solely with β-arrestin without G-protein involvement, and undergoing internalization thereby regulating chemokine availability ^38^.

To analyze GPR15LG-mediated receptor activation, we used the NanoBit technology to monitor β-arrestin-2 recruitment. In this assay, receptor-β-arrestin-2 interactions are detected by luminescence generated when one enzyme subunit, fused to the C-termini of GPR15, CXCR4, or ACKR3, comes into proximity with a second enzyme subunit fused to the N-terminus of β-arrestin-2 ^39^ ^40^. As expected ^37^, GPR15LG induced a dose-dependent recruitment of β-arrestin-2 to GPR15 with an EC_50_ value of 400 nM (**Figure 3a**) ^2^. CXCL12 failed to activate GPR15, but led to β-arrestin-2 recruitment to CXCR4 and ACKR3, with EC_50_ values of 30 nM for both (**Figure 3a**) ^23,41^. AMD3100 had no effect on GPR15 and CXCR4 but weakly activated ACKR3 (**Figure 3a**), as reported ^42^. GPR15LG induced dose-dependent activation of ACKR3 at concentrations exceeding 1 µM (**Figure 3a**), suggesting it as a weak ACKR3 agonist. Although, this is in line with our modeling results (**Figure 2**), ACKR3 has a strong propensity for activation ^43^, and was not able to compete with CXCL12 binding to ACKR3. If the ACKR3/GPR15LG interaction plays a biological role remains to be confirmed. In contrast, GPR15LG did not induce β-arrestin-2 recruitment to CXCR4 (**Figure 3a**) or CXCR4 internalization (**Figure 3b-d**). GPR15 and CXCR4 are GPCRs and binding of an agonist leads to G-protein dissociation. In line with this, GPR15LG induced G-protein dissociation at GPR15, but not at CXCR4 or ACKR3 (**Figure 3e**). Consistent with a lack of agonistic function, a CXCR4-expressing murine acute lymphoblastic leukemia (ALL) cell line, which strongly releases Ca^++^ upon stimulation with CXCL12, showed no Ca^++^ release upon GPR15LG treatment (**Figure 3f**).

**Figure 3.**
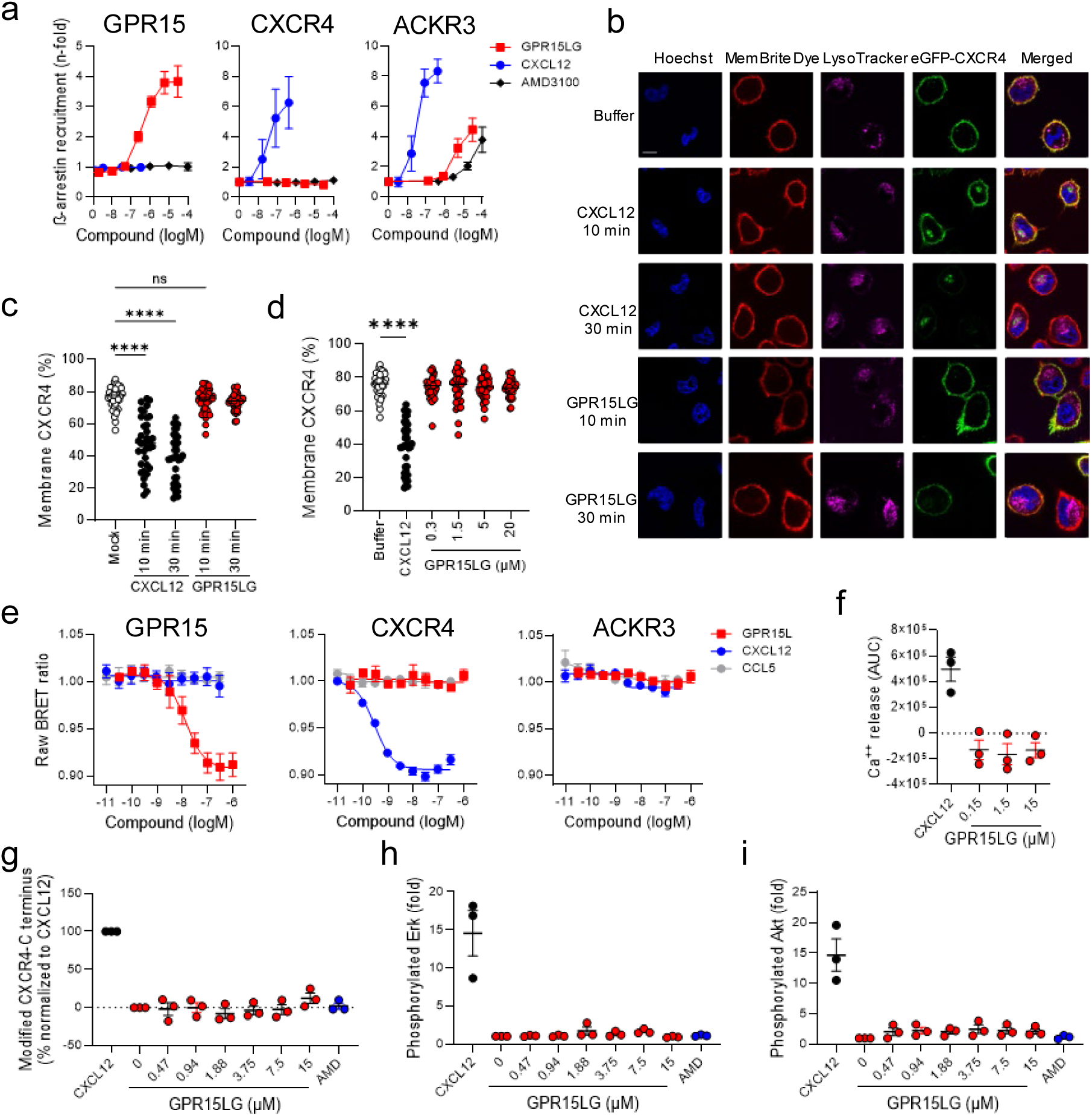
GPR15LG does not induce CXCR4 downstream signaling. a) β-arrestin-2 recruitment to GPR15, ACKR3, and CXCR4. HEK293T cells transiently expressing the respective GPCRs and β-arrestin-2 were treated with increasing concentrations of ligands. After 1 hour, fold changes of areas under the curves (AUC) were calculated. Confocal microscopy (b) and quantification (c, d) of HeLa cells transiently expressing eGFP-CXCR4 (green). Cells were treated with 30 nM CXCL12 or 20 µM GPR15LG for 10 or 30 minutes or a serial dilution for 30 min and stained with Hoechst (blue) for nuclei, MemBrite Dye (red) for cell membrane, and LysoTracker (pink) for lysosomes. Scale bar = 7.5 µm. Data represent mean of *n = 35* individual cells. e) Gαi-protein dissociation was performed in HEK293T cells using a NanoBRET assay. f) Quantification of Ca^+^⁺ release from BCR-ABL1-transformed mouse B cells treated with mouse CXCL12 or indicated concentrations of GPR15LG. Shown are the AUC for each measurement. (g, h, i) C-terminal modifications or phosphorylation of CXCR4, Akt, or Erk in primary CD4⁺ T cells. Cells were stimulated with serially diluted GPR15LG and stained for: (g) anti-CXCR4 mAb (UMB2) to detect unmodified C-terminal epitopes, (h) anti-pErk mAb, and (i) anti-pAkt mAb. a, e-i) Data represent mean ± SEM, *n = 3*, performed in triplicates. **** = p < 0.0001, ns = not significant (One-way ANOVA with Dunnett multiple comparison test).

So far, experiments were performed in cells overexpressing CXCR4. To assess activation of CXCR4 downstream pathways in relevant immune cell subtypes, we isolated primary CD4^+^ T cells from human blood. These cells showed low GPR15 surface expression and no detectable expression of ACKR3 but were strongly positive for CXCR4 as determined by flow cytometry (**Figure S5**). Cells were stimulated with increasing concentrations of GPR15LG and analyzed for modification of the receptor C-terminus and phosphorylation of Akt and Erk. In contrast to CXCL12, which modified the CXCR4 C-terminus (**Figure 3g**) and activated Akt and Erk upon binding (**Figure 3h, i**), GPR15LG did not activate any of these pathways (**Figure 3g-i**). Thus, GPR15LG is an agonist for GPR15 and a weak agonist for ACKR3, but not for CXCR4.

### 2.3. GPR15LG acts as a biased antagonist for CXCR4

We next tested, if GPR15LG binding to CXCR4 interferes with CXCL12-induced signaling. Preincubation with both GPR15LG or AMD3100 dose-dependently inhibited CXCL12-induced β-arrestin-2 recruitment to CXCR4 with IC_50_ values of 3 µM and 0.2 µM, respectively (**Figure 4a**). Accordingly, pretreatment with GPR15LG also inhibited CXCL12-induced CXCR4 internalization (**Figure 4b, c**).

**Figure 4.**
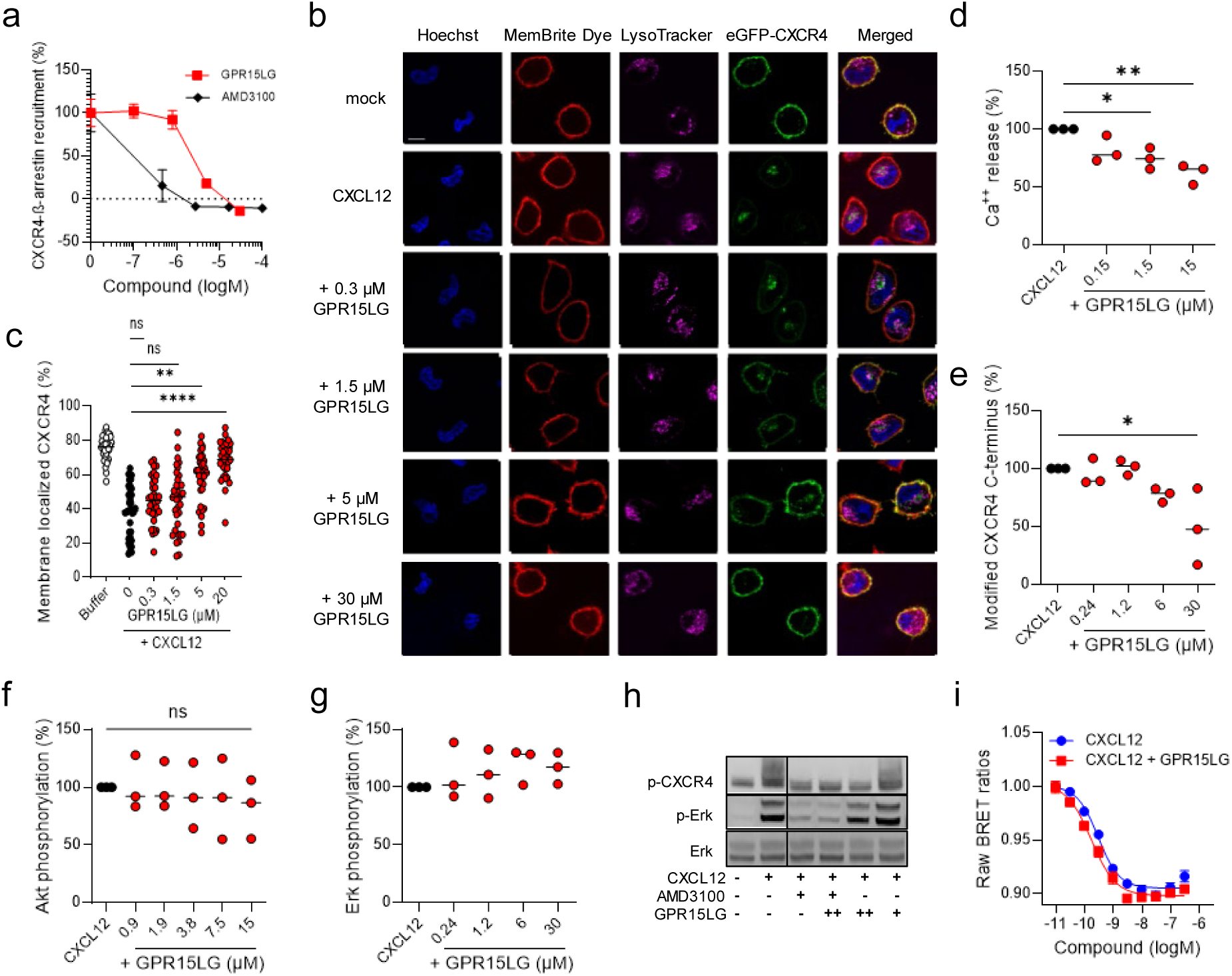
GPR15LG antagonism is biased towards the β-arrestin-2 signaling pathway. a) Inhibition of CXCL12-induced β-arrestin-2 recruitment to CXCR4 by GPR15LG and AMD3100 in HEK293T cells. b) Confocal microscopy images (b) and quantification (c) of HeLa cells transiently expressing eGFP-CXCR4 (green). Cells were pre-treated with serial dilutions of GPR15LG for 30 minutes, followed by CXCL12 treatment for 30 minutes. Cells were stained with Hoechst (blue, nuclei), MemBrite Dye (red, cell membrane), and LysoTracker (pink, lysosomes). Scale bar = 7.5 µm. Data represent mean of *n = 35* individual cells. d) Quantification of Ca^+^⁺ release from BCR-ABL1-transformed mouse B cells treated with CXCL12 with or without GPR15LG. Areas under the curve (AUCs) for each Ca²⁺ measurement were calculated and normalized to the CXCL12 control. e) Inhibition of CXCL12-induced modification of the CXCR4 C-terminus by GPR15LG, measured by flow cytometry staining. f, g) Effect of GPR15LG on CXCL12-induced Akt (f) or Erk (g) phosphorylation in primary CD4⁺ T cells. a, d-g) Data are shown as mean ± SEM, *n = 3*, performed in triplicates. h) Western blot analysis of cells stimulated with CXCL12 in the presence of GPR15LG. One representative image from three independent experiments is shown. AMD3100 = 100 mM, GPR15LG = 1.5 µM (+) or 30 µM (++). i) Gαi-protein dissociation induced by CXCL12 in the presence or absence of 300 nM GPR15LG was performed in HEK293T cells using a NanoBRET assay. *p < 0.1, **p < 0.01, ****p < 0.0001 (One-way ANOVA with Dunnett multiple comparison test).

At higher concentrations, pretreatment with GPR15LG reduced CXCL12-indcued Ca^++^ signaling in the BCR-ABL ALL cell line (**Figure 4d**), and attenuated CXCL12-mediated CXCR4 C-terminal modification in primary CD4^+^ T cells (**Figure 4e**). Nonetheless, GPR15LG had no effect on CXCL12-induced CXCR4 C-terminus, Akt and Erk phosphorylation in primary CD4^+^ T cells (**Figure 4f-h**) or blocked CXCL12-driven G-protein dissociation (**Figure 4i**). These findings suggest that GPR15LG acts as a biased antagonist for CXCR4, specifically targeting β-arrestin-2 recruitment and receptor internalization while sparing other CXCR4 signaling pathways. Of note, GPR15LG had no antagonistic effect on β-arrestin-2 recruitment towards a panel of other chemokine receptors (CX3CR1, CCR6, CCR7, CCR2, CXCR1, and CCR5), indicating specificity for GPR15, ACKR3 and CXCR4 (**Figure S6**).

### 2.4. GPR15LG synergistically enhances CXCL12-induced cell signaling and migration

GPR15LG has been reported to promote chemotaxis of GPR15-expressing effector T cells ^2^. However, the involvement of receptors beyond GPR15 remained unclear. To explore whether GPR15LG binding to CXCR4 modulates CXCL12-induced migration, we performed a transwell migration assay.

As expected, CXCL12 induced dose-dependent migration of primary CD4^+^ T cells, with highest rates observed at concentrations above 100 ng/ml (**Figure 5a**). In contrast, GPR15LG alone did not promote migration on its own. Notably, at concentrations below 320 nM, GPR15LG enhanced CXCL12-mediated migration. This effect was most notable at low CXCL12 concentrations, where migration was weak but significantly enhanced by GPR15LG (**Figures 5b, c**). For instance, 2.4 nM CXC12 led to a weak specific CD4^+^ T cells migration rate of 6 %. In the presence of 320 nM GPR15LG it was increased to 36 % (**Figure 5b**).

**Figure 5.**
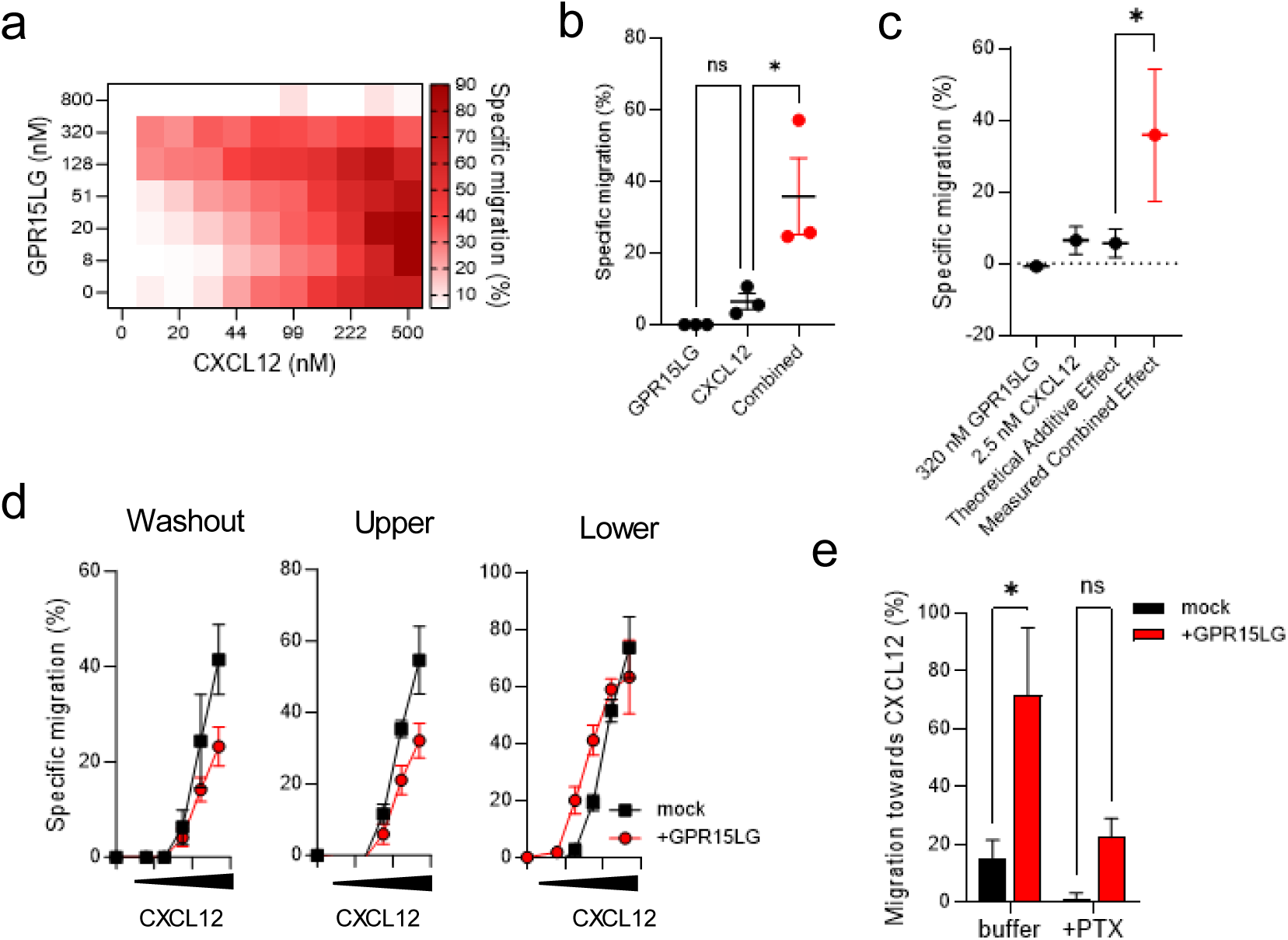
GPR15LG synergistically enhances CXCL12-directed migration of primary CD4^+^ T cells. a) Migration of primary CD4⁺ T cells towards CXCL12 and GPR15LG gradients. GPR15LG and CXCL12 were titrated in a checkerboard format, and cells were allowed to migrate for 4 hours. Data represent the mean of, *n= 3* performed in singlets. b) Specific migration of primary CD4⁺ T cells towards 320 nM GPR15LG, 2.4 nM CXCL12, and their combination. c) Synergism calculation for the combined effect of GPR15LG and CXCL12 migration from panel (a). d) Migration of primary CD4⁺ T cells after different treatment conditions. Cells were pretreated with 100 nM GPR15LG, washed, and then allowed to migrate towards CXCL12 (washout), or incubated with GPR15LG in the upper chamber and allowed to migrate towards CXCL12 (lower), or allowed to migrate towards both CXCL12 and GPR15LG (lower) for 4 hours. e) Migration of primary CD4^+^ T cells towards CXCL12 with and without GPR15LG in the presence of 1 µg/ml Pertussis toxin. b-e) Data are presented as mean ± SEM, *n = 3*, performed in triplicates. *p < 0.1 (b,c: One-way ANOVA with Tukey multiple comparison test; e: Two-Way ANOVA).

To quantify the interaction between GPR15LG and CXCL12, we calculated the Bliss synergy score (S_Bliss_). Assuming independent effects of GPR15LG and CXCL12, the theoretical additive effect was calculated and compared to the measured effects. The S_Bliss_ is then defined as the difference between measured and theoretical additive effects. Values of 0 indicate no interaction, while negative values indicate antagonism and positive values synergistic effects ^44^. When considering all concentrations, the S_Bliss_ was -7.44 ± 4.05 (SD), reflecting an inhibitory effect of GPR15LG at high concentrations (>320 nM). However, when focusing on GPR15LG concentrations below 320 nM, where enhancement occurred, the S_Bliss_ reached values exceeding 20, indicating strong synergy between GPR15LG and CXCL12 in promoting cell migration (**Figure S7**).

Interestingly, the synergistic effects observed in the migration assay appeared to contradict our earlier findings, where GPR15LG either antagonized CXCL12 or had no effect. To resolve this, we investigated whether the timing and spatial presentation of GPR15LG affected its activity. Migration assays were conducted under three conditions: (i) cells preincubated with GPR15LG and washed before migration, (ii) GPR15LG in the upper chamber and CXCL12 in the lower chamber, and (iii) both GPR15LG and CXCL12 present together in the lower chamber. As expected, GPR15LG inhibited CXCL12-mediated migration in the preincubation and upper chamber conditions. However, when GPR15LG and CXCL12 were co-present in the lower chamber, migration was enhanced, suggesting that GPR15LG must directly interact with CXCL12 to exert a synergistic effect (**Figure 5d**).

Notably, both, CXCL12-induced migration and GPR15LG-mediated enhancement were sensitive to pertussis toxin, indicating a G-protein-independent mechanism (**Figure 5e**).

To assess whether GPR15LG cooperates with CXCL12 in enhancing CXCR4 downstream pathways, we tested conditions where both chemokines were added simultaneously. When diluted in a checkerboard titration, incubated for 30 minutes, and used to stimulate primary CD4^+^ T cells, GPR15LG enhanced CXCL12-induced Akt and Erk phosphorylation (**Figure 6a-d**). This effect was most pronounced at low CXCL12 concentrations, with pAkt increased 3-fold and pErk rose 2.5-fold at 7.4 nM CXCL12 and 600 nM GPR15LG (**Figures 6b, d**). GPR15LG did not inhibit β-arrestin-2 recruitment to CXCR4 when premixed with CXCL12 and even slightly enhanced CXCL12-induced recruitment (**Figures 6e, f**). However, GPR15LG continued to antagonize CXCL12-induced CXCR4-eGFP internalization when both ligands were premixed (**Figure 6g**). Furthermore, GPR15LG enhanced β-arrestin-2 recruitment to ACKR3 in the presence of CXCL12 (**Figures 6h, i**), while CXCL12 had no effect on GPR15 recruitment (**Figure S8**). Thus, GPR15LG increases CXCL12-induced CXCR4 and ACKR3 signaling, particularly for G-protein-dependent pathways and chemotaxis.

**Figure 6.**
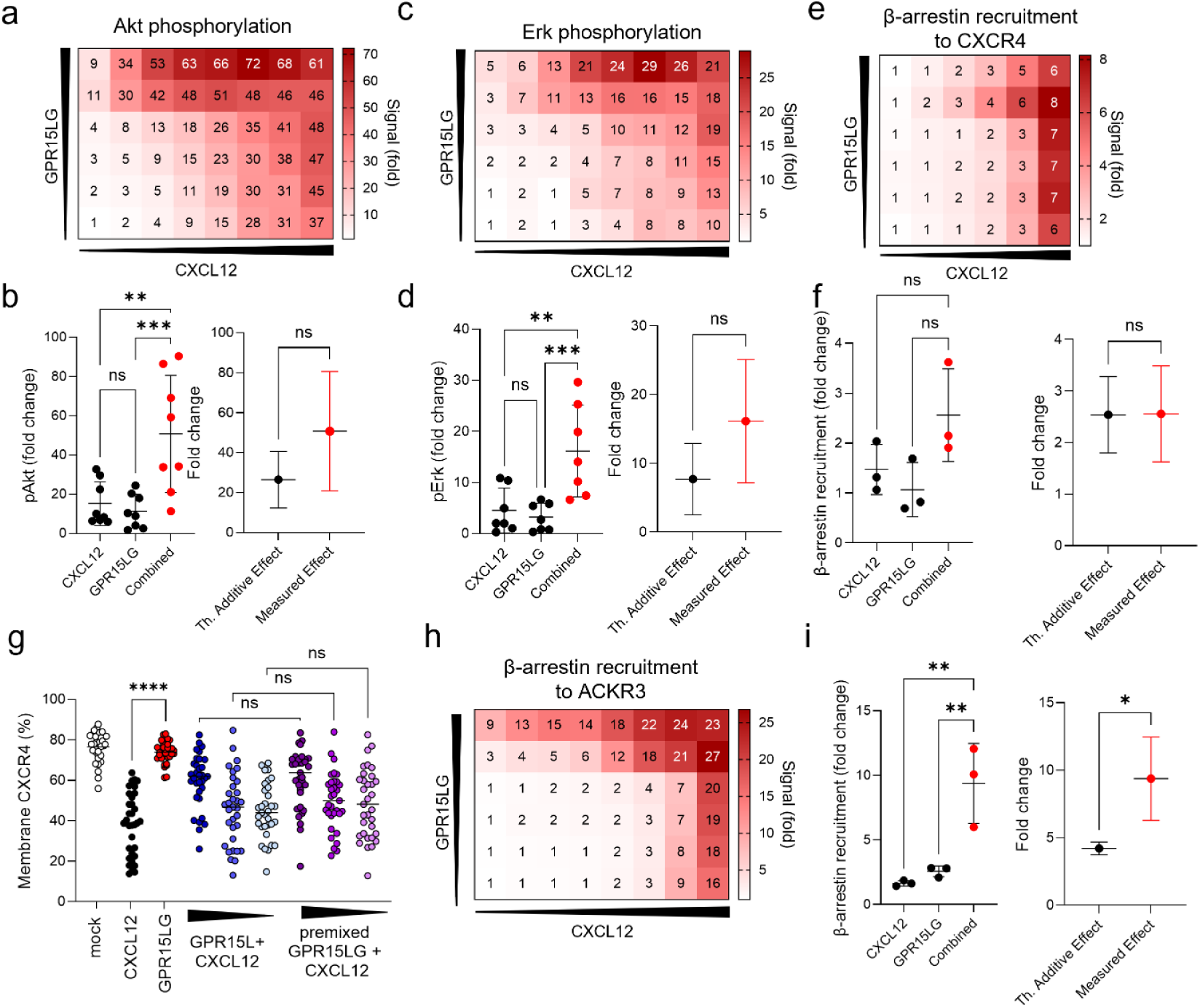
GPR15LG enhances CXCL12-induced downstream signaling, but not receptor desensitization. a-d) Primary CD4⁺ T cells were treated with serial dilutions of GPR15LG (1:5, starting at 3 µM) and CXCL12 (1:3, starting at 200 nM) for 30 minutes at 37°C in PBS. Phosphorylation of Akt (a,b) and Erk (c,d) was assessed by flow cytometry 2 minutes post-treatment. b,d) Assessment of the synergism for the combination of 600 nM GPR15LG and 7.4 nM CXCL12 for panels (a) and (c). e) Recruitment of β-arrestin-2 to CXCR4 by combined CXCL12 and GPR15LG. f) Assessment of the synergism for the combination of 600 nM GPR15LG and 7.4 nM CXCL12 from panel (e). g) Quantification of membrane CXCR4-eGFP expression in HeLa cells. Cells were treated with CXCL12, GPR15LG, or pretreated with GPR15LG for 30 minutes before CXCL12 was added, or treated with both GPR15LG and CXCL12 in combination. Data represent the mean of *n = 35* individual cells. h, i) Recruitment of β-arrestin-2 to ACKR3 by combined CXCL12 and GPR15LG. Data for synergism are presented in panel (i). a-c, h) Data are shown as mean, *n = 3-8*, performed in singlets. *p < 0.1, **p < 0.01, ****p < 0.0001 (b,d,f,i: One-way ANOVA with Tukeys multiple comparison test or unpaired t-test for synergism. g: One-Way ANOVA with Sidak multiple comparison test).

### 2.5. Synergistic effects of GPR15LG and CXCL12 on immune and cancer cell migration

Migration and signaling assays were conducted using primary CD4^+^ T cells, which express high levels of CXCR4 but low levels of GPR15 and ACKR3 (**Figure S5**). To investigate whether GPR15LG synergizes with CXCL12 to enhance migration across a broader range of immune cell subtypes, we extended our analysis to peripheral blood mononuclear cells (PBMCs), which represent a diverse population of immune cells. We used either a low concentration of CXCL12, which on itself only induces minimal migration, 300 nM GPR15LG, or both in the lower chamber of transwell assays. Migration was significantly enhanced when both chemokines were present, and this effect was blocked by the CXCR4 antagonist AMD3100, indicating a CXCR4-dependent mechanism (**Figure 7a, Figure S9**). Flow cytometry analysis of migrating PBMC subsets showed that the proportions of CD4^+^ T cells, CD8^+^ T cells, and NK cells remained consistent across conditions, indicating that GPR15LG broadly enhances CXCL12-mediated migration rather than targeting specific immune subsets (**Figure 7b, Figure S10**). Notably, GPR15LG alone weakly attracted CD4^+^ T cells but strongly enhanced migration of CD56^+^ NK cells, despite the absence of detectable GPR15 expression on NK cells (**Figure 7b, Figure S5, Figure S10**). This suggests potential involvement of another chemotactic receptor.

**Figure 7.**
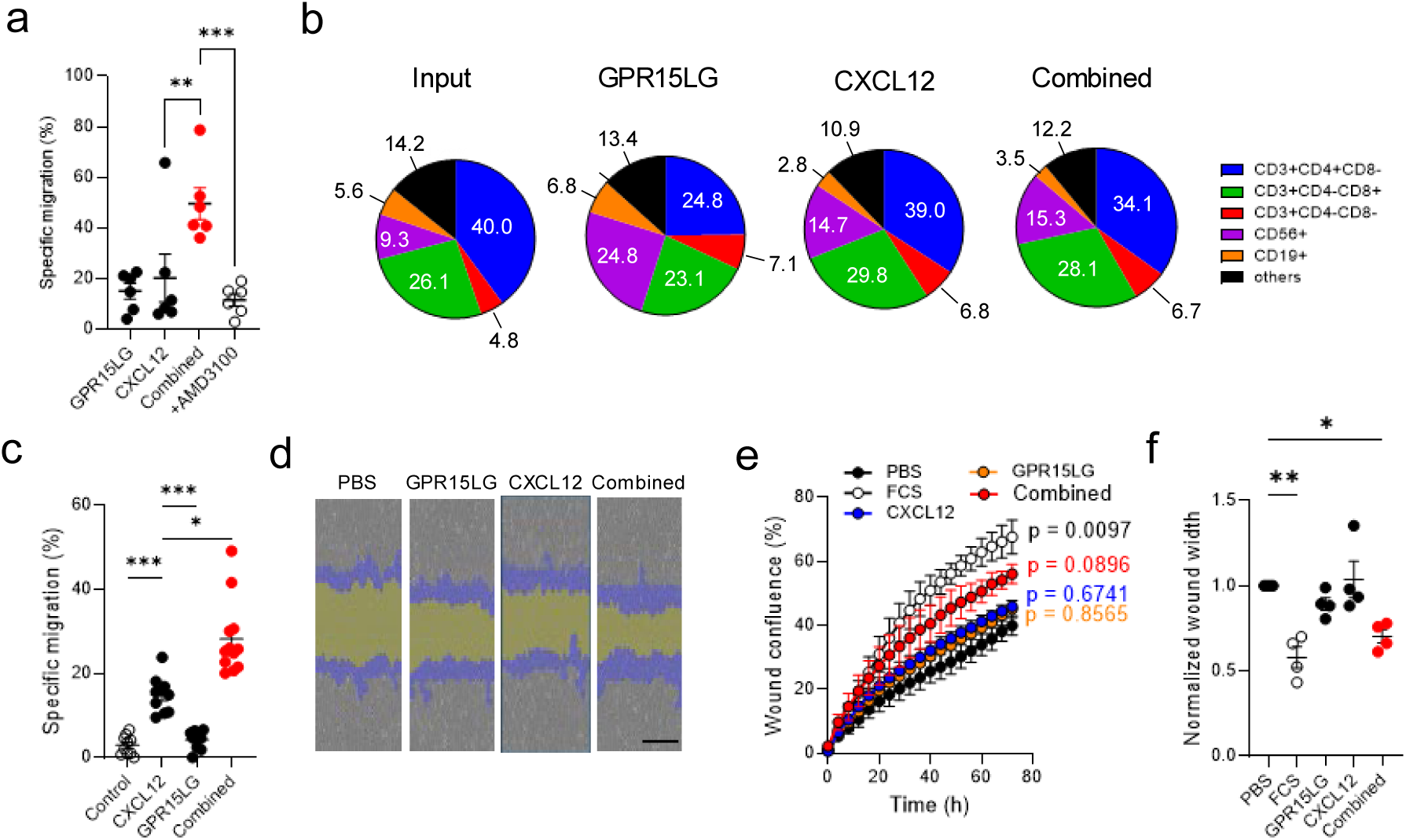
GPR15LG synergistically enhances CXCL12-mediated migration of different immune cell populations and cancer cells. a-b) Migration of healthy peripheral blood mononuclear cells (PBMCs) in the presence of 300 nM GPR15LG, 1.2 nM CXCL12, or their combination with or without 100 µM AMD3100. a) Relative migration was measured using CellTiterGlo after 4 hours of incubation. Data represent mean ± SD, *n = 6 donors*. b) Relative population frequencies of migrated PBMCs. Subsets were identified by flow cytometry, and data are presented as percentages of live single cells. c) Migration of BCWM.1 cells in the presence of 0.5 µM GPR15LG, 60 nM CXCL12, or their combination. d) Representative images from the scratch wound healing assay in HeLa cells at 10x magnification. Remaining “wound width” (yellow) and cells which have grown into the wound (“wound confluence” (blue)) are indicated. e, f) Wound closure in HeLa cells was monitored using the Incucyte Scratch Wound Analysis. After washing, cells were treated with respective ligands which had been incubated for 30 minutes at 37°C in medium before being added to the wells. Wound closure was monitored every 4 hours for 72 hours. Data represent mean ± SEM, *n = 3-4*, performed in triplicates. * p ≤ 0.1, ** p ≤ 0.01, *** p ≤ 0.001, **** p ≤ 0.0001 (one-way ANOVA with Tukey multiple comparison test).

CXCL12 is a major chemoattractant in B cell malignancies, including Waldenström’s macroglobulinemia (WM) ^45^. Despite expression of GPR15 (**Figure S11**), GPR15LG did not directly attract WM cells. However, GPR15LG significantly enhanced CXCL12-directed migration, suggesting a potential role in cancer cell migration and disease progression (**Figure 7c**). The effects of CXCL12 and GPR15LG on solid tissue migration, particularly in wound healing, were also examined. A scratch assay was performed on CXCR4-expressing HeLa cells, where CXCL12 and GPR15LG alone had minimal effects on cell migration. However, their combination significantly increased migration into the scratch, resulting in a smaller wound area (**Figure 7d-f, Figure S12**).

These results demonstrate that GPR15LG enhances CXCL12-induced migration across various cell types in a CXCR4-dependent manner, with this effect being blocked by AMD3100.

## 3. Discussion

Our study reveals that GPR15LG, beyond its established role as a GPR15 ligand, interacts with CXCR4 and weakly ACKR3, impacting CXCL12-driven signaling pathways and cell migration. A key finding is the synergistic effect of GPR15LG with CXCL12, enhancing CXCL12-mediated migration and signaling in both, immune and cancer cells. The CXCR4-specific antagonist AMD3100 blocked this effect, confirming its CXCR4 dependency. These synergistic effects may have significant implications in physiological and pathological contexts. On the one hand, the enhancement of CXCL12 signaling by GPR15LG could contribute to improved adaptive immune responses and accelerated wound healing. On the other, this synergy might also facilitate the spread of cancer cells, underscoring its potential role in tumor progression and metastasis. Understanding these dual effects is critical for evaluating GPR15LG as a therapeutic target or modulator in different disease settings.

GPR15LG shares structural similarity with the CC-chemokine family, although it is not classified as a classical chemokine ^3^. While classical chemokines primarily signal through their N-terminal domains, GPR15LG interacts and activates GPR15 via its C-terminal domain ^46,47^. In contrast, our study demonstrates that GPR15LG engages the CXCR4 binding pocket through conserved residues at its N-terminus, a binding mode that allows it to modulate CXCR4 without fully activating it. Pretreatment with GPR15LG blocked CXCL12 binding to CXCR4 and inhibited β-arrestin-2-dependent downstream signaling, while sparing G-protein-mediated pathways, suggesting a biased antagonistic or allosteric effect on CXCR4. Of note, we acknowledge that the use of different experimental systems and cell types might limit conclusions drawn in our study. More detailed analyses are needed to confirm the biased effects of GPR15LG towards CXCR4. Dual-functions mediated by the C- and N-termini have been described for other chemokines. For example, peptides derived from the CCL21 C-terminus are allosteric enhancers of CCL21-induced migration, whereas the N-terminus is responsible for receptor activation ^48^. This highlights that modulation of classical chemokine-receptor interactions by other cytokines or their peptides might be a common phenomenon.

Our data show that GPR15LG is an antagonist for CXCR4 at high concentrations, and synergistically enhances CXCL12-mediated functions at lower concentrations. Dose-dependent differences in chemokine function have been observed before. For example, at low concentrations, CXCL12 attracts a variety of CXCR4^+^ leukocyte subpopulations. However, at high concentrations CXCL12 induces a repulsive effect, indicating a dose-dependent regulatory mechanism preventing excessive T cell infiltration ^49^. Thus, a similar regulatory mechanism could down-modulate GPR15LG and CXCL12 induced cell migration, when concentrations reach a critical level.

Synergistic interactions with CXCL12 have been observed for other chemokines and non-chemokine proteins ^50–52^. The High Mobility Group Box 1 (HMGB1), a stress-associated molecule, enhances CXCL12’s chemotactic potential by forming a heterocomplex with it ^50^. Our data suggest that GPR15LG might employ a similar mechanism by staying in close proximity to CXCL12 to enhance migration. However, further studies are needed to confirm whether a heterocomplex is formed and if this complex interacts with CXCR4 ^51,52^. Recent studies have shown that the orphan chemokine CXCL14 enhances chemotaxis of cells expressing CXCR4, CXCR5 and CCR7 ^51,52^. Although the mechanism remained unclear, the authors suggested that CXCL14 enhances receptor oligomerization thereby reducing activation thresholds. It would be interesting to see, if GPR15LG has similar effects after binding to CXCR4. GPR15LG could also enhance CXCL12 interaction with glycosaminoglycans (GAGs) on the cell surface, or mobilize CXCL12 from GAGs thereby enhancing local concentrations^53^. Alternatively, other receptors, such as ACKR3, Sushi Domain Containing 2 receptor (SUSD2) ^14^, or Mas-related G protein-coupled receptors (MRGPRs) ^54^, might be involved.

GPR15LG’s enhancement of CXCL12-induced migration was observed across various immune cell populations, including CD8^+^ T cells, CD4^+^ T cells, B cells, and NK cells. This effect might exacerbate inflammation in conditions where CXCL12 is upregulated. Studies have identified GPR15LG as pruritogen or driver of inflammation in the skin. However, these effects were mediated by a GPR15-independent mechanism ^11,54,55^: Tseng et al. showed that GPR15LG causes GPR15-independent mast cell degranulation and stimulation of neuronal cells thereby causing an itch response ^54^. In another study, GPR15LG induced a pro-inflammatory response in GPR15-negative keratinocytes and affected skin barrier formation ^11^. In addition, Sezin et al. have demonstrated that GPR15LG aggravates psoriasiform dermatitis in GPR15-deficient mice ^55^. It is tempting to speculate that GPR15LG-mediates pro-inflammatory effects by modulating the CXCR4/CXCL12 axis. In the context of wound healing, GPR15LG has been shown to promote granulation tissue formation, collagen deposition, and angiogenesis ^56^. Our observation that GPR15LG enhances CXCL12-induced wound closure suggests that these effects are mediated via CXCR4.

The observed synergistic effect offers promising therapeutic opportunities. In diseases where excessive CXCR4 signaling exacerbates pathology, such as certain inflammatory and metastatic cancers ^18^, targeting GPR15LG could attenuate CXCL12-driven responses. GPR15LG-based modulators or antagonists might ameliorate skin inflammation and metastatic spread in cancers. Future studies should explore the molecular basis of the GPR15LG-CXCL12 synergy through structural analyses. Studies on GPR15LG’s impact on various immune cell subsets and cancer cell types to clarify its broader role in CXCR4 signaling and its potential as a therapeutic target are highly warranted. Finally, investigating the effects of GPR15LG on skin manifestations of B cell or T cell lymphomas and its potential role in wound healing will provide valuable insights into its diverse functions.

## 4. Conclusion

In summary, GPR15LG emerges as a novel and versatile modulator of the CXCR4/CXCL12 axis, with context-dependent effects that range from synergy to biased antagonism. These findings underscore the complexity of chemokine receptor signaling and highlight the potential of GPR15LG as a therapeutic target in inflammatory and oncologic diseases. Future research should focus on the structural basis of GPR15LG’s interactions with CXCR4 and other receptors, its potential impact on CXCL12-GAG dynamics, and its effects *in vivo* to fully understand its diverse roles and therapeutic applications.

## 5. Materials and Methods

### Reagents and GPR15LG synthesis

GPR15LG was synthesized by Bachem (Bubendorf, Switzerland). All other chemokines were obtained from PeproTech (Rocky Hill, NJ, USA). AMD3100 was purchased from Sigma Aldrich.

### Cloning of β-arrestin-2 reporter constructs

The previously published CXCR4-LgBiT, ACKR3-LgBiT and SmBiT-arrestin plasmids were generously donated by Jong-Ik Hwang (Department of Biomedical Sciences, Korea University, Seoul) ^57^. GPR15-LgBiT was assembled by Gibson assembly, utilizing the previously published pcDNA3.1_GPR15 ^17^ and the commercially available N196 pBiT1.1-C [TK/LgBiT] Vectors (Promega, N2014). PCR with Phire II Hot Start Polymerase (Thermo Fisher Scientific) was performed on both plasmids with their respective primers (biomers.net, Ulm, Germany). Subsequent Gibson assembly of the PCR products using the NEBuilder® HiFi DNA Assembly Master Mix and sequencing revealed an unwanted insertion of an additional Cysteine at the C-terminus of the GPR15 sequence which was repaired by Q5 Site-directed mutagenesis (New England Biolabs, E0554S) with the indicated primers (Table 1).

**Table 1.**
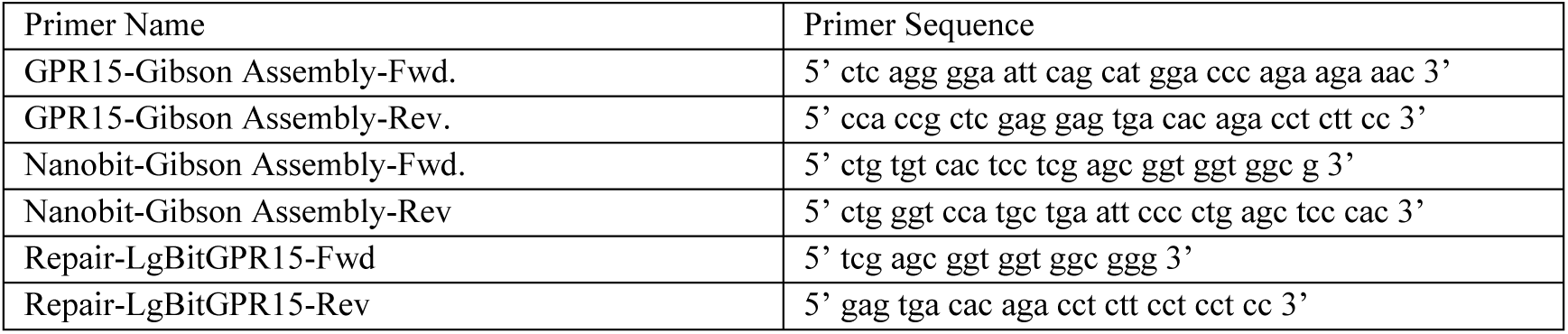
Primer used for β-arrestin-2 reporter constructs.

All other chemokine receptor-LgBiT constructs were generated by de-novo synthesis at Twist Bioscience (South San Francisco, California, USA). In short, the mRNA coding sequences for each receptor (Table 2) were accessed at the NIH genebank, linked to a 16-AA Ser-Gly linker and then to the pBiT1.1-N[TK/LgBiT] expression sequence, available from Promega (Rocky Hill, NJ, USA). These sequences were then inserted into the pTwist EF1 Alpha vector and synthesized de-novo. Plasmids were amplified with the Wizard Plus Midiprep DNA Purification System (Promega) according to the manufacturer’s instructions.

**Table 2.**
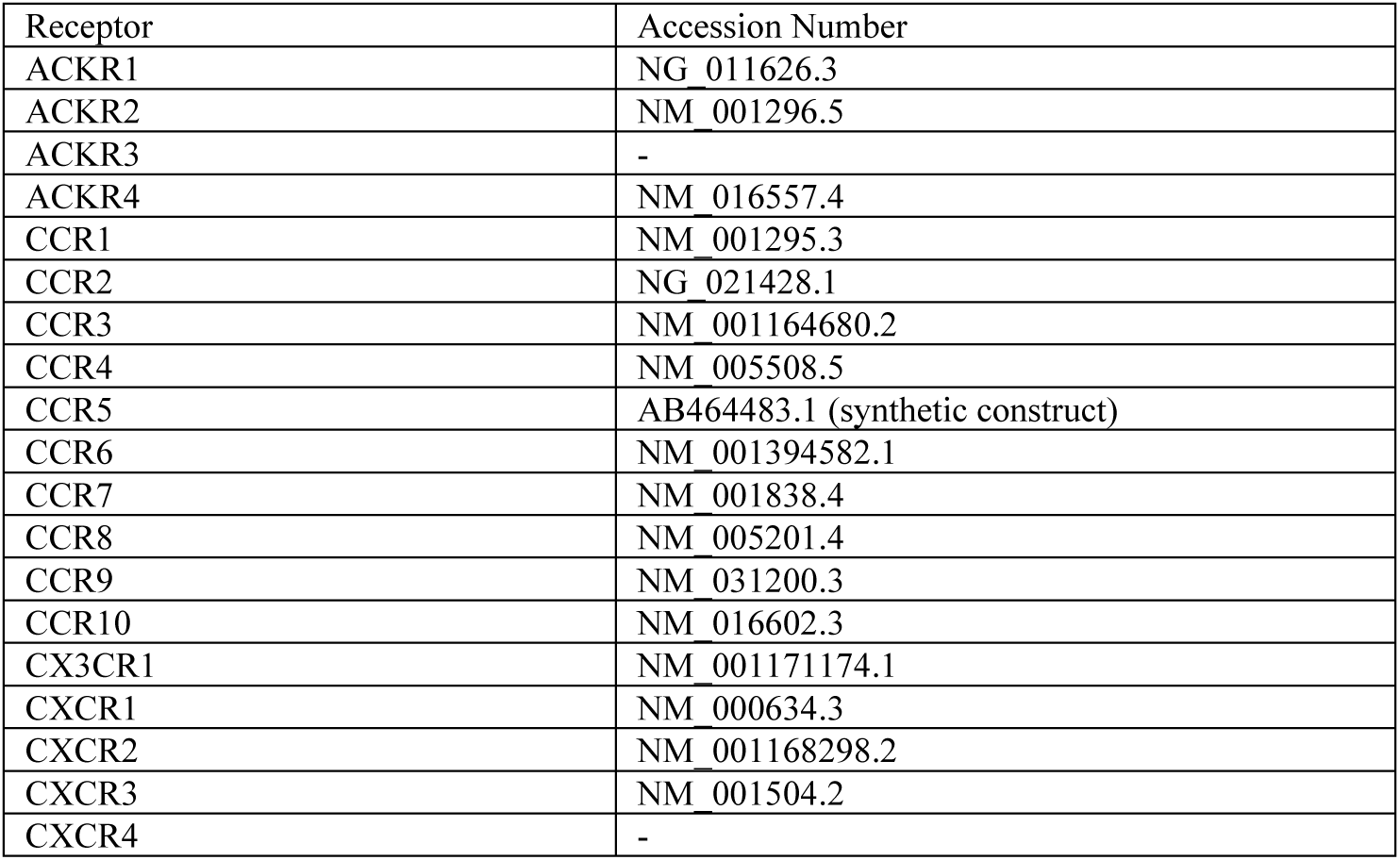
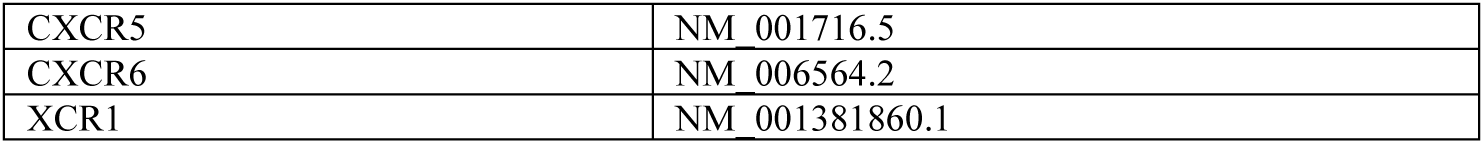
Accession number of chemokine receptors used for β-arrestin-2 recruitment assays.

| Receptor | Accession Number |
| --- | --- |
| ACKR1 | NG_011626.3 |
| ACKR2 | NM_001296.5 |
| ACKR3 | - |
| ACKR4 | NM_016557.4 |
| CCR1 | NM_001295.3 |
| CCR2 | NG_021428.1 |
| CCR3 | NM_001164680.2 |
| CCR4 | NM_005508.5 |
| CCR5 | AB464483.1 (synthetic construct) |
| CCR6 | NM_001394582.1 |
| CCR7 | NM_001838.4 |
| CCR8 | NM_005201.4 |
| CCR9 | NM_031200.3 |
| CCR10 | NM_016602.3 |
| CX3CR1 | NM_001171174.1 |
| CXCR1 | NM_000634.3 |
| CXCR2 | NM_001168298.2 |
| CXCR3 | NM_001504.2 |
| CXCR4 | - |
| CXCR5 | NM_001716.5 |
| CXCR6 | NM_006564.2 |
| XCR1 | NM_001381860.1 |

### Site-directed CXCR4 mutagenesis and cloning experiments

Site directed mutagenesis was performed as previously published ^35^. In short: Q5 Site-directed mutagenesis (New England Biolabs, E0554S) utilizing commercially synthesized primers (biomers.net, Ulm, Germany) (Table 3) was used to introduce different point mutations (E179A, D187A) in the previously published pcDNA3.1_hCXCR4_GRP construct. Subsequently, a second and third site-directed mutagenesis was carried out. (E179A plus D181A; E179A plus D181A and D182A; E179A plus D193A and E268A; D187A plus D181A and E288A; D179A plus D193A and D262A) All primers are listed in the table below (Table 3). Final constructs were sequenced before usage.

**Table 3.**
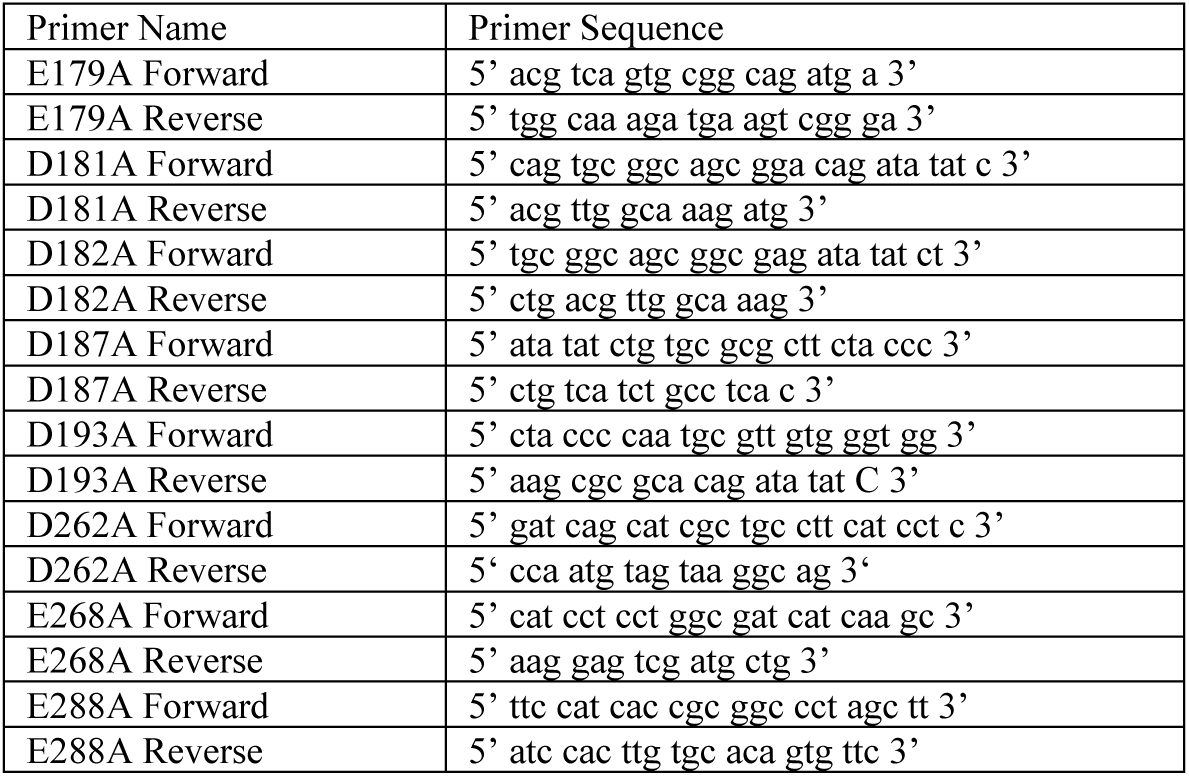
Primer used for CXCR4 mutations.

### Cell culture

HEK293T, SupT1 cells and HeLa cells were obtained from the American Type Culture Collection (ATCC). BCWM.1 cells (Cellosaurus RID: CVCL_A035) expressed BCR subtype IgM, λ, derived from Waldenström’s macroglobulinemia patient’s bone marrow ^58^. TZM-bl cells were obtained the NIH AIDS Reagent Program. Primary human PBMCs were obtained from healthy donors by Ficoll density centrifugation. CD4^+^ T-cells were isolated from buffy coats with the RosetteSep CD4^+^ T-cell enrichment cocktail according to the manufacturer’s instruction followed by Ficoll density centrifugation. For all experiments, CD4^+^ T-cells were treated overnight with 10 ng/ml Interleukin-2 from Miltenyi Biotec (Bergisch Gladbach, Germany). TZM-bl and HEK293T cells were maintained in Dulbecco’s modified Eagle’s medium (DMEM) supplemented with 10% fetal calf serum (FCS), 100 units/mL penicillin, 100 µg/ml streptomycin and 2 mmol/L L-glutamine. SupT1, PBMC and CD4^+^ T-cells cells were maintained in Roswell Park Memorial Institute medium (RPMI) supplemented with 10% or 1% FCS, 100 units/mL penicillin, 100 µg/ml streptomycin, 2 mmol/L L-glutamine and 1 mmol/L HEPES (Gibco). BCR-ABL1-transformed ALL cells were generated in house by retroviral transduction of BCR-ABL1 construct into mouse bone marrow derived pre-B cells ^59^. The cells were cultured in Iscove’s Modified Dulbecco’s Media (Sigma) supplemented with 10% heat-inactivated fetal bovine serum (FBS, PAN Biotech), 2 mM L-glutamine (Gibco), 100 units/ml of penicillin/streptomycin (Gibco) and 50 μM beta-mercaptoethanol (Gibco) at 37°C in a 7.5% CO_2_ incubator.

### Staining of cell surface receptors

PBMCs were washed with PBS once and stained with NucFix Viability Dye (ThermoFischer) in PBS for 20 min at room temperature. Then, cells were stained for pan population markers and surface GPCR expression using human α-CD3-Alexa Fluor 700, α-CD8-PerCP-Cy5.5, α-CD56-BV650, α-CD19-BV421, α-CD4-BV605, α-CXCR4-APC-Cy7, α-GPR15-PE/Dazzle594 and α-CXCR7-PE (all Biolegend) in BD Horizon Brilliant Stain Buffer (Invitrogen), with 10% FcR Blocking Reagent (Miltenyi Biotec) for 1 hour at 4°C. After staining, cells were washed with FACS buffer once and fixed with 2% PFA before performing flow cytometry.

### Competition with CXCR4-antibodies

Competition with the CXCR4 antibodies was performed on SupT1 cells. Compounds were serially diluted in cold PBS and afterwards added to 5 × 10^4^ cells. APC-conjugated anti-human CXCR4 antibody targeting ECL-2 (clone 12G5, #555976, BD) or PE-conjugated anti-human CXCR4 antibody targeting the N-terminus (clone 1D9, #551510, BD) were diluted in PBS containing 1% FCS and added immediately afterwards. After 90 min incubation at 4 °C, unbound antibody was removed by washing twice in PBS containing 1% FCS, cells were fixed in 2% paraformaldehyde and analyzed by flow cytometry. Mean fluorescence intensity of isotype staining was subtracted from all sample MFIs and values were normalized to full stain (100%) and unstained (0%).

### CXCR4 and ACKR3 ligand binding competition monitored by NanoBRET

Ligand binding to CXCR4 and ACKR3 was monitored by NanoBRET. Briefly, HEK293T cells stably expressing the receptor N-terminally fused to Nanoluciferase were distributed into white 96-well plates (5 × 10^4^ cells per well). Increasing concentrations of ligands were added to the cells, as well as CXCL12_AZ568_ (10 nM) and incubated for 2 hours on ice. The Nanoluciferase substrate was then added and donor emission (450/8 nm BP filter) and acceptor emission (600 nm LP filter) were immediately measured on a GloMax Discover plate reader (Promega). BRET binding signal was defined as acceptor/donor ratio, and cells not treated with CXCL12_AZ568_ were used to define 0 % BRET binding, whereas cells that were treated with CXCL12_AZ568_ alone were used to define 100 % BRET binding.

### HIV-1 production and inhibition

Viral stocks of CXCR4-tropic and CCR5-tropic HIV-1 were generated by transient transfection of HEK293T cells with proviral DNA as described before ^60^. Inhibition of viral infection was performed in TZM-bl reporter cells. For this, 1 × 10^4^ cells (in growth medium supplemented with 2.5% FCS) were pretreated with serially diluted inhibitors for 30 min at 37 °C. Cells were then inoculated with virus diluted in serum-free media. Infection rates were determined after 3 days using Gal-Screen system (Applied Biosystems). Virus controls without compound were set as 100% for normalization.

### CXCL12 induced Akt, Erk phosphorylation and modification of the CXCR4 C-terminus

CXCL12-induced phosphorylation of Erk, Akt and the CXCR4 C-terminus were monitored by phosphoflow cytometry. For this 100,000 CD4^+^ T cells were starved overnight at 37 °C (RPMI medium supplemented with 1% FCS). Inhibitors were then added for 10 min and cells subsequently stimulated with compounds for 2 min. The reaction was stopped by adding 2% paraformaldehyde and shifting to 4 °C for at least 10 min. Cells were then permeabilized with ice cold methanol and stained with phospho-p44/42 MAPK (ERK1) (Tyr204)/(ERK2) (Tyr187) (D1H6G) mouse mAb (Cell Signaling, #5726), phospho-Akt (Ser473) (193H12) Rabbit mAb #4058 or the CXCR4 rabbit mAb (UMB2) and adequate secondary antibodies for flow cytometry. UMB2 targets the unmodified C-terminus of CXCR4 no longer binds upon phosphorylation. Average mean fluorescence (MFI) of the unstained control was subtracted from all values to avoid autofluorescence and unspecific binding and values were normalized (100% = Average CXCL12 MFI, 0%: Average buffer only MFI).

### Ca^++^ signaling

Ca^++^ signaling was analyzed as described previously ^34,59^. Briefly, BCR-ABL1 transformed mouse B cells were collected as 1×10^6^ cells/treatment and loaded with Calcium-sensitive dye Indo-1 AM (#I1223, Invitrogen) and 0.5 mg/mL of pluronic F-127 (#P3000MP, Invitrogen) in respective media supplemented with 1% FBS at 37°C for 45 min. Cells were then washed and treated with the inhibitors for 10 min at 37°C. Baseline signal for calcium was measured for 30sec by flow cytometry followed by stimulation with 100 ng/mL of mouse CXCL12 or different concentrations of GPR15LG. The area under the curve (AUC) of each calcium flux plot was determined using FlowJo (version 10). The AUC of water control (solvent for CXCL12) was subtracted from each treatment to get the correct estimation of Calcium signal upon CXCL12 stimulation.

### G protein dissociation assay

The measurement of Gαi protein dissociation upon CXCR4, GPR15 and ACKR3 activation was performed using NanoBRET-based assay. HEK-293T cells were plated in a 6-well plate (0.8 x 106 per well), cultured for 24h before transfection with a polycistronic vector encoding Gα subunit of G proteins fused to the Nanoluciferase and the Gβγ dimer-fused to a circular permutated Venus fluorescent protein ^61^. 24 hours after transfection, cells were harvested, incubated for 3 minutes at 37°C with Nano-Glo Live Cell substrate, and distributed into white 96-well plates, each well containing 1.5 x 105 cells. Ligands were then added and BRET signal was measured with a GloMax plate reader (Promega) equipped with 450/8 filter for donor luminescence emission and 530 LP filter for acceptor fluorescence emission. BRET signal was defined as acceptor/donor ratio.

### β-arrestin recruitment assay

For transfection, 20.000 HEK293T cells were plated out into 96-well Eppendorf microplates with reservoirs (Eppendorf Hamburg, Germany) (cat. 0030 730.135). One day later, cells were transiently transfected with 50 ng of GPCR-LgBiT plasmid and 50 ng of SmBiT-Arrestin plasmid, using TransIT-LT1®, purchased from Mirus Bio (Madison, WI, USA) according to the manufacturer’s instruction in Opti-MEM. After an additional 24 hours, for antagonistic measurement, all medium was replaced with 100 µl of Opti-MEM and reservoirs in the microplate were filled with PBS. Plates were incubated at 37°C for 30 min to stabilize temperature before 25 µl of Nano-Glo Live Cell Reagent was added to each well and an initial baseline luminescence was recorded for 10 min at a Synergy H1. Subsequently 15 µl of serially diluted inhibitory compound or buffer control were added for an additional 10 min. Finally, 10 µl of stimulatory compounds were added and the kinetic was run for 1 hour. For agonistic measurements, medium was changed to 110 µl of Opti-MEM and after addition of 25 µl of Nano-Glo Live Reagent for 10 min, 15 µl of serially diluted stimulatory ligand or buffer control were added for 1 hour. Baseline values were averaged and fold change to average baseline was calculated. Fold changes were then normalized to average buffer fold change for each timepoint and area under the curve was calculated. For normalization of antagonistic measurement, area under the curve (AUC) values of buffer were set to 0%, values of stimulatory compound were set to 100%. For agonistic measurements fold change to buffer control was calculated. β-arrestin-2 recruitment screens on additional GPCRs were performed as described above, with changes to the volumes: 55 µl (antagonism) or 65 µl (agonism) of Opti-MEM were used with final volumes in the well at 100 µl.

### Chemotaxis

Chemotaxis assays of CD4^+^ T-cells were performed in 96-well transwell assay plates (Merck Milipore, Carrigtwohill, Ireland), while assays with PBMCs were performed in 24-well or 96-well transwell assay plates (Corning) with 5 μm polycarbonate filters. Plates were blocked with a 1% BSA solution for 30 min at room temperature before cell seeding to prevent unspecific binding of chemokines to plastic. Then 7.5 × 10^4^ cells in 50 μL (96-well) or 4 × 10^5^ in 100 µL (24-well) assay buffer (RPMI supplemented with 0.1% bovine serum albumin) were seeded into the upper chambers. 170 μL (96-well) or 600 µL (24-well) assay buffer supplemented with compounds or input control cells were filled into bottom chambers. Cells were allowed to migrate towards ligands by combining upper and lower chambers for 4 hours at 37 °C (5% CO_2_). Relative migration was measured for each donor by CellTiterGlo® assay (Promega, Madison, WI, USA). Specific migration was calculated by normalizing luminescence values to input control (100%) and buffer control (0%). Migrated cells in the lower chamber were collected and stained for surface marker and GPCR expression and analyzed via flow cytometry. Migration of BCWM.1 cells was performed in a 96 well transwell plate system with an 8.0-µM pore size (Corning). BCWM.1 cells were resuspended in serum-free RPMI media and 100,000 cells in 50 µL volume were seeded into the upper chamber of the plate. The lower chamber of the plate was filled with 150 µL of serum-free RPMI media with or without chemokines. Cells that migrated into the lower chamber were harvested after 4 hours of incubation at 37° C, 5% CO_2_, and counted.

### Scratch assay

Scratch Assays were performed following the Incucyte Scratch Wound Assay (Sartorius, Germany) protocol. Hela cells were seeded at 15.000/cells per well of a 96-well Imagelock Plate (Sartorius, Germany). After 24 hours a wound was generated simultaneously in each well using the Incucyte 96-Well Woundmaker Tool (Sartorius, Germany). After washing twice with PBS, wells were treated with negative control (no FCS), positive control (+FCS), GPR15LG (0-3600nM), CXCL12 (0-100ng/ml) or the combination of GPR15LG and CXCL12. The ligands were incubated for 30 min at 37°C degree before treatment. The plate was imaged using an Incucyte Live-Cell Analysis System every 4 hours for 72 hours. Images of the wound were acquired using 10x magnification and the “Scratch Wound Analysis Software Module”. Analysis of “wound width” and “wound confluence” were performed with the background/cell separation set to “1”. “Wound width” was normalized to the negative control of each experiment.

### Receptor internalization

6 × 10^4^ HeLa cells were plated on 13mm borosilicate cover slips in 24-well cell culture plates in 500 µL medium. After 24h incubation cells were transfected with CXCR4-eGFP fusion protein, using TransIT-LT1®, 500 ng of DNA per well in 50 µl Opti-MEM with 1,5 µL LT-1. After 6 h of incubation at 37°, medium was exchanged to remove transfection reagents. 48h post-transfection, cells were treated with 200 µl of medium containing respective concentrations of ligands for 10 or 30 minutes at 37°C. Medium was then exchanged for pre-warmed medium containing Hoechst 33342 Solution at 1:2000 for 30 min at 37°C. Cells were washed three times in PBS and fresh pre-warmed medium with 75 nM Lyso Tracker Deep Red L12492 was added for 30 min at 37°C. Cells were washed again in PBS three times and MemBrite® Fix Pre-Staining Solution 1X in PBS was added for 5 minutes at RT. Cold PBS with MemBrite Fix Dye 30095 diluted 1:1000 was then added for 30 min at 4°C. Cells were again washed three times in ice-cold PBS and fixed with 4% PFA for 15 min at RT. Cells were again washed three times in PBS and once in distilled water to remove salts. Coverslips were mounted on glass slides using 10 µl of Mowiol Mounting Medium and dried at 4°C overnight. Images were acquired using a LSM 710 system by Zeiss (Oberkochen, Germany), 35 cells per condition. Signal intensity of CXCR4 at cell surface and in the cytoplasm was quantified using the open-source software ImageJ.

### Molecular modeling

Biomolecular simulations were conducted to investigate the binding and dynamic behavior of the CXCR4/GPR15LG and ACKR3/GPR15LG complexes using Gaussian accelerated Molecular Dynamics (GaMD) ^62^ to enhance conformational sampling. The CHARMM36m force field^63^, as implemented in AMBER 20 ^64^, was employed. The CXCR4 model derived from prior studies ^35^ (comprising sequence 1-319) served as the foundation for constructing the complete 3D structure of CXCR4. To achieve this, the missing C-terminal segment (320-352) was taken from a model predicted using AlphaFold2 ^65^ and integrated within the previously reported model ^35^ to produce the full-length CXCR4. The 3D structures of ACKR3 (AF-P25106) and GPR15LG (AF-Q6UWK7) were obtained from the AlphaFold Protein Structure Database (accessed on https://alphafold.ebi.ac.uk/) ^66,67^.

The initial geometries of the complexes (GPR15LG/CXCR4, GPR15LG/ACKR3) were generated by placing GPR15LG 100 Å away from the center of mass of CXCR4 or ACKR3 in three different orientations (Table 4). The systems were embedded in a lipid bilayer of POPC molecules and subsequently placed in a solvent box of explicit water molecules. The TIP3P water model ^68^ was used and NaCl was added for neutralization. The CHARMM-GUI membrane builder ^69,70^ was employed for setting up all systems and generating their topologies for CHARMM36m ^71^.

**Table 4.** Simulation setups and their composition.

| **CXCR4** with | Box dimensions | Number of POPC molecules | Total number of atoms |
| --- | --- | --- | --- |
| GPR15LG (pose1) | 170 Å x 170 Å x 254 Å | 713 | 600425 |
| GPR15LG (pose2) | 150 Å x 150 Å x 260 Å | 538 | 467868 |
| GPR15LG (pose3) | 160 Å x 160 Å x 238 Å | 623 | 502637 |
| **ACKR3** with | Box dimensions | Number of POPC molecules | Total number of atoms |
| GPR15LG (pose1) | 150 Å x 150 Å x 244 Å | 539 | 406115 |
| GPR15LG (pose2) | 150 Å x 150 Å x 238 Å | 539 | 397460 |
| GPR15LG (pose3) | 160 Å x 160 Å x 226 Å | 623 | 438560 |

Each system was subjected to 5000 minimization steps, followed by six equilibration runs of 1875 ps in total as implemented in CHARMM-GUI, applying harmonic positional restraints at 303.15 K and 1 bar of external pressure. Then, nine ns of classical MD were carried out, followed by 40 ns of GaMD equilibration. Unrestrained production GaMD runs (3 replicas of 400 ns each) with a time step of 2 fs were performed for all set-ups. The last 200 ns of each replica of each pose were concatenated, resulting in 1800 ns (3 x 3 x 200 ns) of simulation trajectory all systems. All the frames were aligned onto the transmembrane domain (TMD, residues 30 – 300) of CXCR4 and (TMD, residues 40-310) of ACKR3. The clustering analysis of the backbone of the GPR15LG protein was performed using a Visual Molecular Dynamics (VMD) plug-in (https://github.com/luisico/clustering), with an RMSD cutoff of 23 Å. The centroids of the clusters were identified using the VMD plug-in considering the average position of all the atoms in the trajectory frames within the cluster. These structures allowed identifying the most populated poses of the protein-protein complexes. We analyzed the key interactions that contribute to the stabilization of each complex by using the hydrogen bond plug-in of VMD with a distance cutoff between heavy atoms of 3.5 Å and an angle cutoff of 30 degrees.

### Statistics and sequence alignments

One- or two-way ANOVA followed by Dunnett’s, Tukey’s or Sidak’s multiple comparison tests were performed as indicated using GraphPad Prism version 10.3.1 for Windows, GraphPad Software, Boston, Massachusetts USA, www.graphpad.com. EC_50_ and IC_50_ values were determined using nonlinear regression. Dose-response curves were fitted using the “[Inhibitor] vs. Normalized response” or “[Agonist] vs. response” equations with variable slope (four parameters). The software calculated EC_50_ and IC_50_ values as the concentration of agonist or inhibitor, respectively, that provoked a response halfway between the baseline and maximum response. The Hill slope was not constrained, allowing the model to determine the steepness of the dose-response curve from the experimental data. Synergism was calculated based on the Bliss independence model ^44^ using the SynergyFinder ^72^ online set as: Readout = Inhibition; Outlier detection = Yes; Curve Fitting = LL4; Synergy Calculation with Bliss; Correction = On. Bliss score synergism of fold-change data was calculated by addition of observed effects and their respective variances and testing for difference to measured effects by t-test. The multiple sequence alignment of GPR15LG was performed using Clustal Omega ^73^. Jalview ^74^ and WebLogo ^75^ were employed for the analysis, visualization and generation of the sequence alignment figures.

### Ethics approval statement

Primary human cells were isolated from blood of healthy donors after informed consent. The work was approved by the local research ethics commission of Ulm University.

## Acknowledgements

We thank Jong-Ik Hwang of the Korea University, Seoul for graciously providing us with the β-arrestin-2 as well as the CXCR4 and ACKR3 NanoGlo constructs.

## Data availability

All raw data and coordinates are available upon reasonable request.

