## Supplement figures for "GPR15LG binds CXCR4 and synergistically modulates CXCL12-induced cell signaling and migration"

Dan Albers et al.

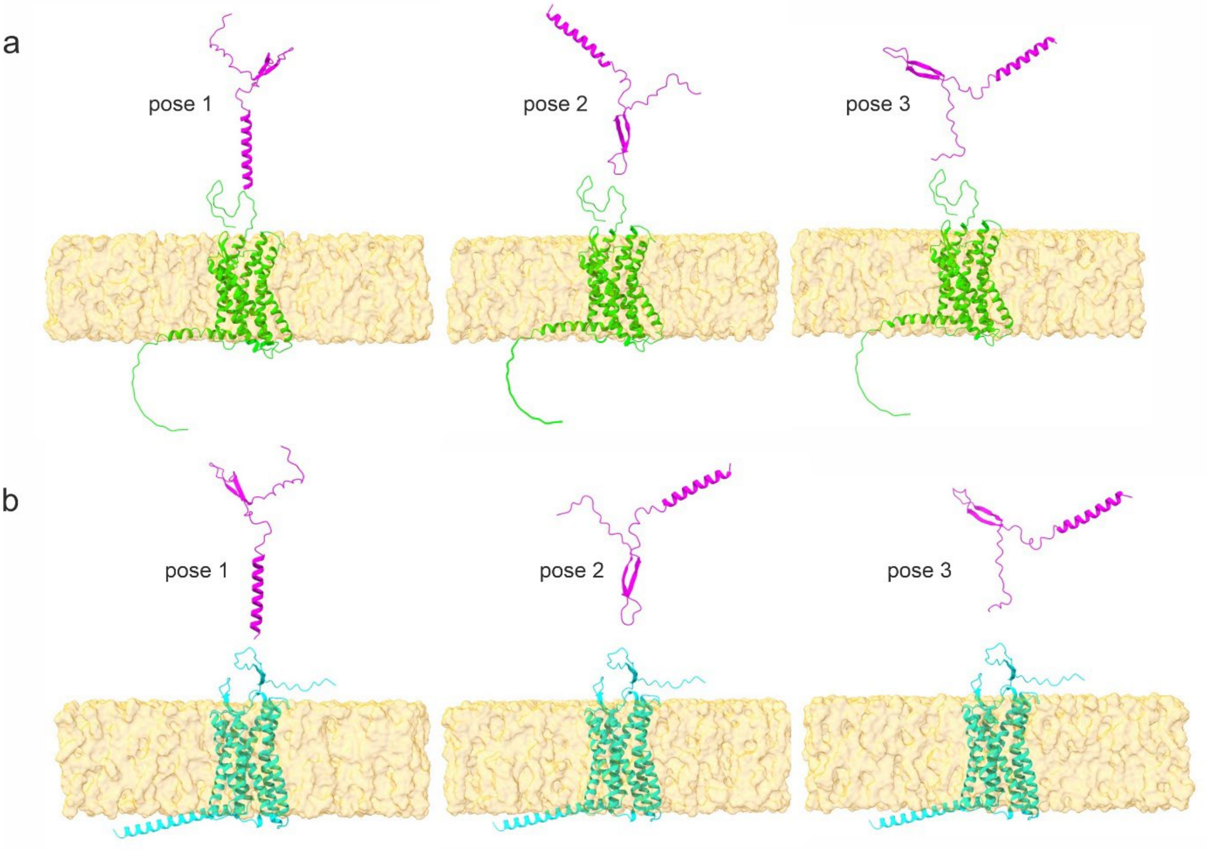


**Figure S1. Initial structures of GPR15LG (magenta) relative to (a) CXCR4 (green) and (b) ACKR3 (cyan) that were employed as starting coordinates for the GaMD simulations.** GPR15LG was placed 100 Å away from the center of mass of the receptors with three different orientations. The POPC membrane is shown as orange surface. Ions and water molecules are not shown for simplicity.

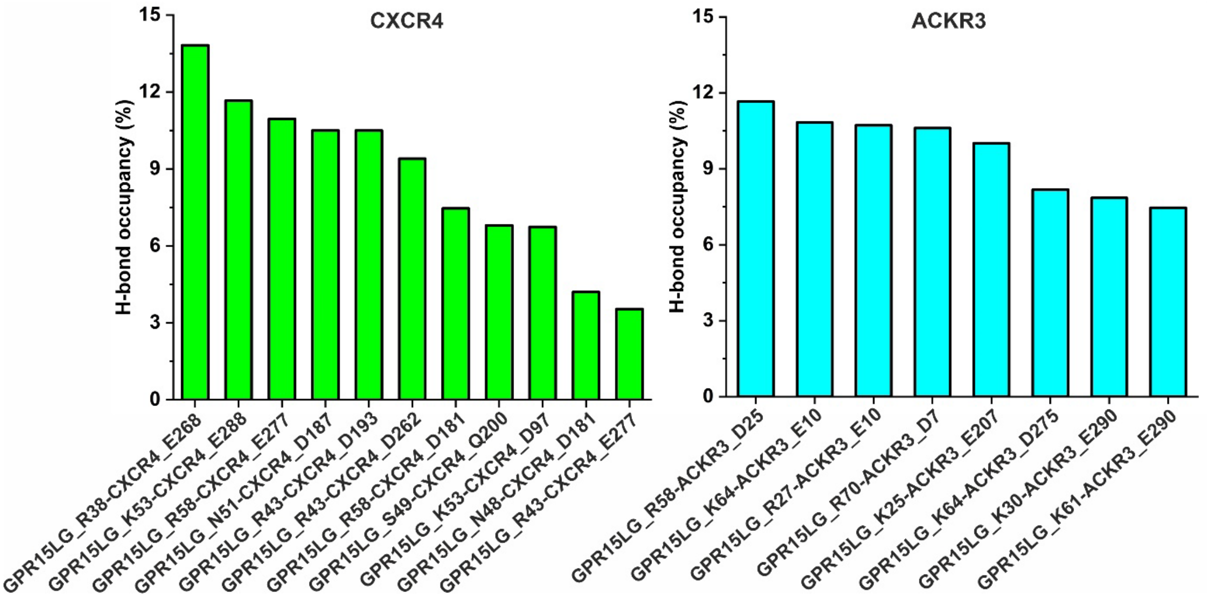


**Figure S2: Occupancy of hydrogen bonds between GPR15LG and CXCR4 (ACKR3) as observed in GaMD simulations.**

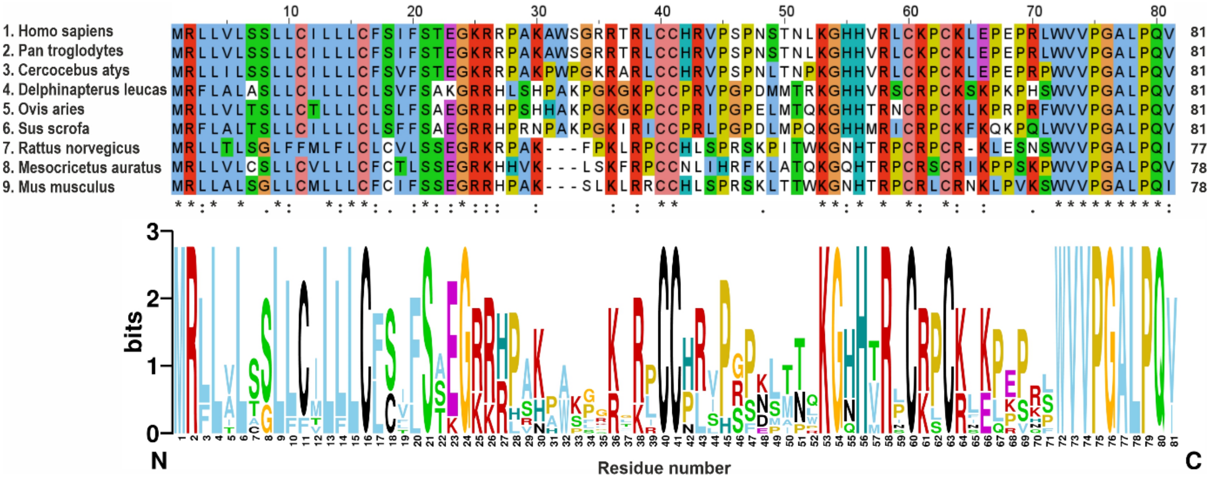


**Figure S3. Multiple sequence alignment and position-based residue conservation of GPR15LG across various species.** Each logo in the bottom panel consists of stacks of symbols for each position in the sequence (generated using Weblogo3: https://weblogo.berkeley.edu/logo.cgi). The total height of a stack indicates the degree of sequence conservation at that position, while the height of individual symbols within the stack denotes the relative frequency of each amino acid at that position.

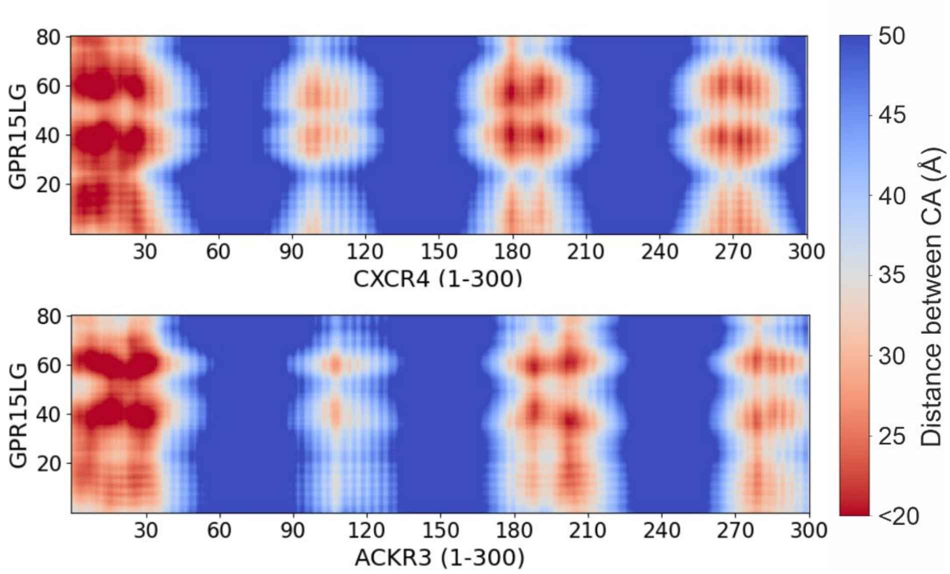


**Figure S4. Maps of the average distance between alpha-carbon (CA) atoms of the full length GPR15LG and the receptors.**

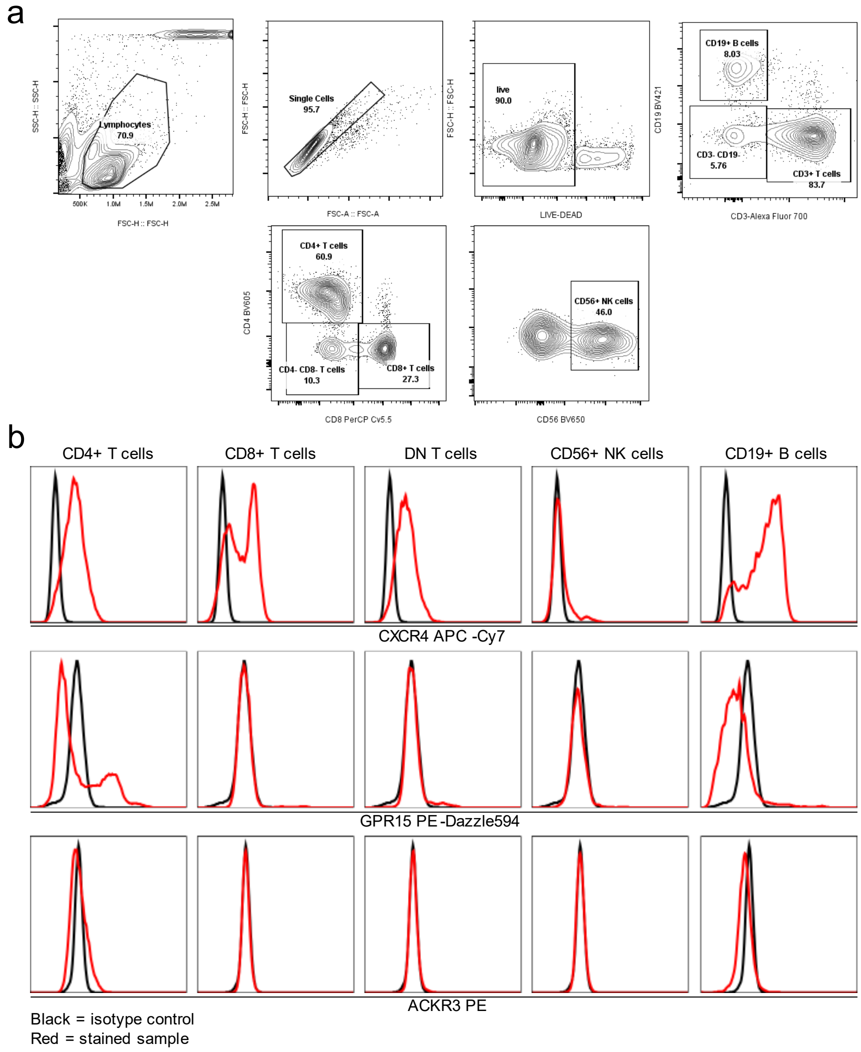


**Figure S5: Surface GPCR expression on main PBMC subsets.** PBMCs were stained for flow cytometry to identify CD19$^{+}$ B cells. CD3$^{+}$ T cells, CD4$^{+}$ and CD8$^{+}$ T cells and natural killer (NK) cells. a) Gating strategy. b) CXCR4, GPR15 and ACKR3 expression on selected PBMC populations. Shown are data from one representative donor.

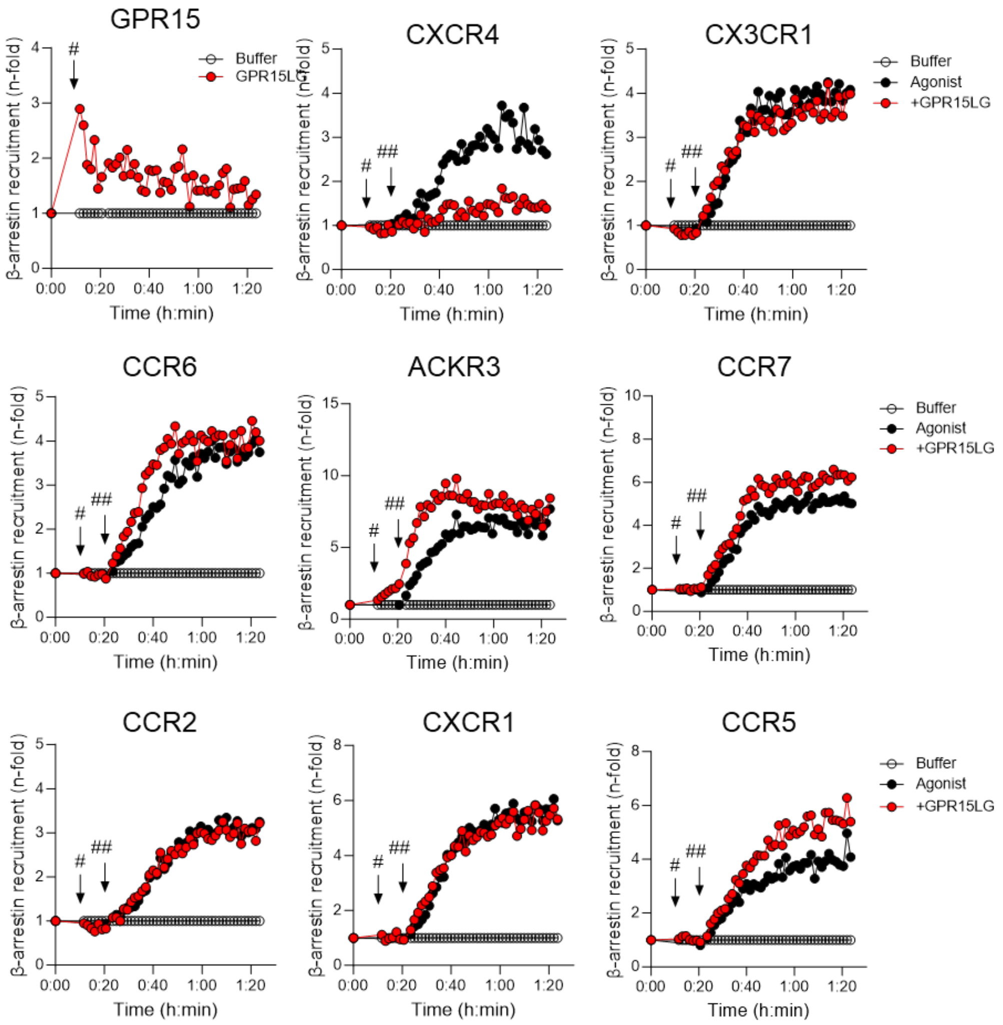


**Figure S6. Effects of GPR15LG treatment on β-arrestin-2-recruitment to a panel of chemokine receptors.** HEK293T cells were transiently transfected with reporter constructs. Baseline was recorded for 10 min, and cells treated with GPR15LG for 10 min (#). Afterwards respective agonists were added (##) and kinetic data were measured for 1 hour. N=1, in duplicates.

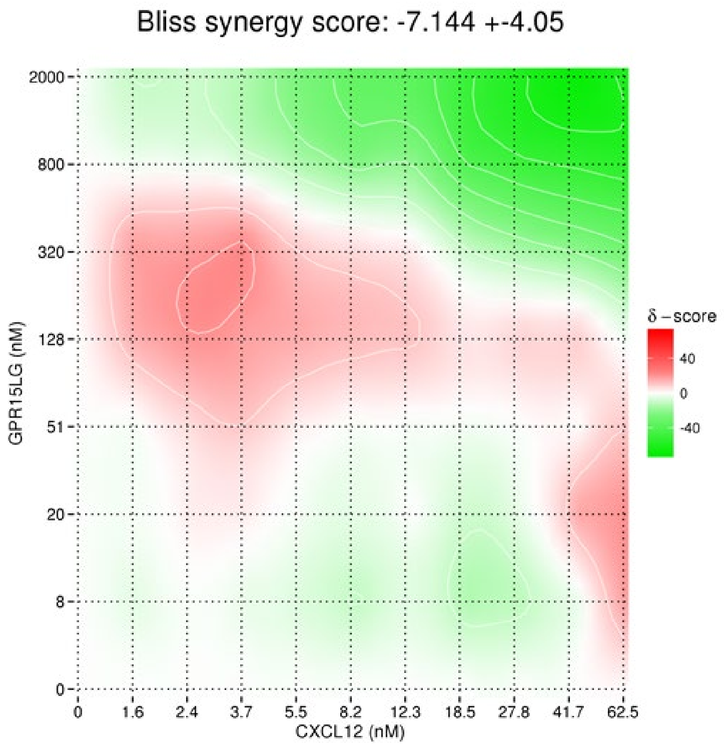


**Figure S7. Quantification of the synergistic effects of GPR15LG on CXCL12-dependent migration.** Migration of $CD4^+$ T Cells was quantified by CellTiterGlo assay and data was evaluated by SynergyFinder. For calculation, data were taken from three independent experiments, performed in singlets.

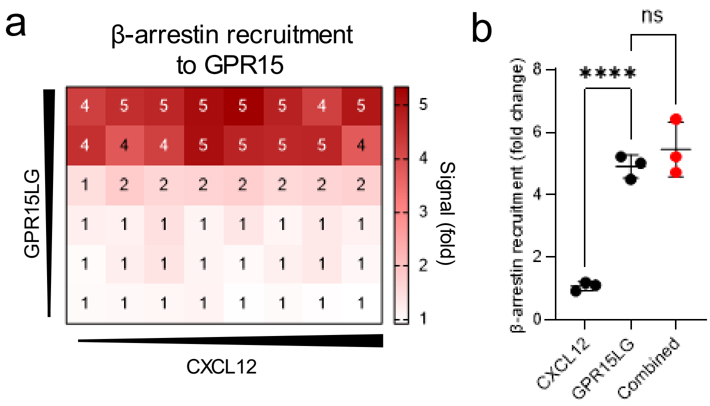


**Figure S8. Recruitment of β-arrestin-2 to GPR15**. a) Recruitment of β-arrestin-2 to GPR15 by combined treatment with CXCL12 and GPR15LG. b) Assessment of the synergism for the combination of 600 nM GPR15LG and 7.4 nM CXCL12 from panel (a). N=3, in singlets ± SD.

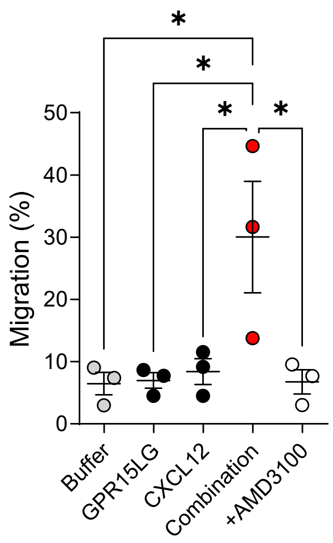


**Figure S9. Migration of PBMCs isolated from older donors to CXCL12 is synergistically enhanced in the presence of GPR15LG.** PBMCs isolated from 3 donors aged 56 to 66 years were seeded in 96 well transwell plates and migration was studied in the presence of CXCL12 (1.2 nM) or GPR15LG (300 nM), or their combination in the presence or absence of 10 µM AMD3100. (One-Way ANOVA with Tukey multiple Comparison test).

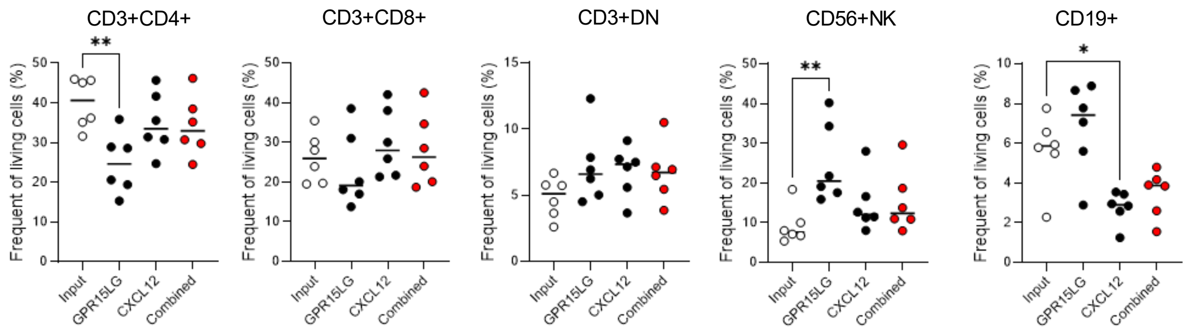


**Figure S10. GPR15LG enhances CXCL12-induced migration independent of the immune cell subset.** Migrated cells were collected and stained for abundance of immune cell subsets in living cells. Subsets were identified using flow cytometry. Shown are data from 6 individual healthy donors. CD3$^+$DN: CD4$^-$ CD8$^-$ CD3$^+$ double negative T cells.

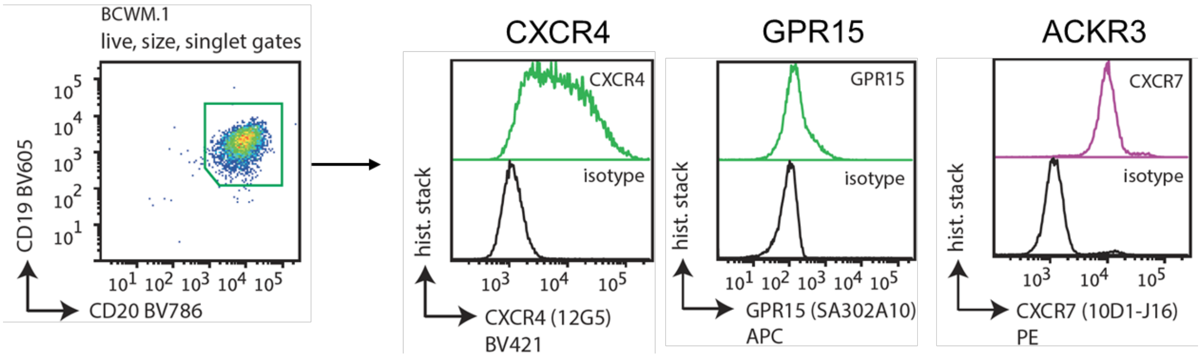


**Figure S11. Cell surface expression of WM cells.**

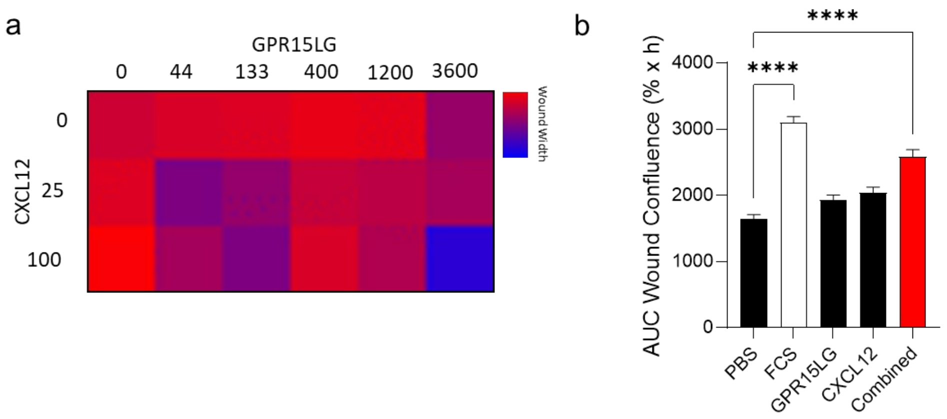


**Figure S12. GPR15LG enhances CXCL12-mediated migration of HeLa cells in a scratch assay.** Scratch assay performed with HeLa cells using the Incucyte Scratch Wound Analysis. After washing, cells were treated with respective ligands which had been incubated for 30 minutes at 37°C in medium before being added to the wells. Wound closure was monitored every 4 hours for 72 hours. a) Heatmap indicating wound width at the respective concentration of either CXCL12, GPR15LG or the mix after 72 hours for one of the three experiments. b) AUC wound confluence of indicated treatments. Data represent mean ± SEM, $n = 3\text{-}4$, performed in triplicates. **** $p \leq 0.0001$ (one-way ANOVA).
